# Spatial and temporal metagenomics of river compartments reveals viral community dynamics in an urban impacted stream

**DOI:** 10.1101/2023.04.04.535500

**Authors:** Josué Rodríguez-Ramos, Angela Oliverio, Mikayla A. Borton, Robert Danczak, Birgit M. Mueller, Hanna Schulz, Jared Ellenbogen, Rory M. Flynn, Rebecca A. Daly, LeAundra Schopflin, Michael Shaffer, Amy Goldman, Joerg Lewandowski, James C. Stegen, Kelly C. Wrighton

## Abstract

Although river ecosystems comprise less than 1% of Earth’s total non-glaciated area, they are critical modulators of microbially and virally orchestrated global biogeochemical cycles. However, most studies either use data that is not spatially resolved or is collected at timepoints that do not reflect the short life cycles of microorganisms. As a result, the relevance of microbiome interactions and the impacts they have over time on biogeochemical cycles are poorly understood. To assess how viral and microbial communities change over time, we sampled surface water and pore water compartments of the wastewater-impacted River Erpe in Germany every 3 hours over a 48-hour period resulting in 32 metagenomes paired to geochemical and metabolite measurements. We reconstructed 6,500 viral and 1,033 microbial genomes and found distinct communities associated with each river compartment. We show that 17% of our vMAGs clustered to viruses from other ecosystems like wastewater treatment plants and rivers. Our results also indicated that 70% of the viral community was persistent in surface waters, whereas only 13% were persistent in the pore waters taken from the hyporheic zone. Finally, we predicted linkages between 73 viral genomes and 38 microbial genomes. These putatively linked hosts included members of the *Competibacteraceae,* which we suggest are potential contributors to carbon and nitrogen cycling. Together, these findings demonstrate that microbial and viral communities in surface waters of this urban river can exist as stable communities along a flowing river; and raise important considerations for ecosystem models attempting to constrain dynamics of river biogeochemical cycles.

## Introduction

Rivers are crucial modulators of global biogeochemical cycles and provide a dynamic, moving passageway between terrestrial and aquatic ecosystems (Allen and Pavelsky, 2018). Corresponding to ∼7% of global CO_2_ and ∼5% of global CH_4_ emissions per year, rivers contribute up to 2,508 Tg yr^-1^of carbon dioxide (CO_2_), and ∼30.5 Tg yr^-1^ of methane (CH_4_) (Villa et al., 2020; Rosentreter et al., 2021; Friedlingstein et al., 2022; Liu et al., 2022). Within rivers, microbial communities are key orchestrators of carbon and nitrogen transformations, where they contribute between 40-90% of total river respiration (Naegeli and Uehlinger, 1997; Battin et al., 2003; Rodríguez-Ramos et al., 2022). Despite a general understanding of the importance of microbial metabolism, knowledge of river viral communities and their impacts on microbial communities remains scarce.

Viruses are the most abundant organism on the planet, with estimates of up to 10^31^ viral particles worldwide (Hendrix et al., 1999; Munn, 2006; Bar-On et al., 2018; Mushegian, 2020). These viral predators are mostly studied in marine ecosystems, where viruses can lyse 20-40% of bacteria daily (Weinbauer, 2004; Weinbauer and Rassoulzadegan, 2004; Suttle, 2007; Chow and Suttle, 2015; Guidi et al., 2016) and play key roles reprogramming their bacterial hosts with ecosystem-wide consequences (Sullivan et al., 2006; Anantharaman et al., 2014; Hurwitz and U’Ren, 2016). Although research has mostly focused on marine ecosystems, recent efforts have been made to expand our knowledge of natural viral communities in freshwater aquatic environments like lakes (Roux et al., 2017; Berg et al., 2021) and estuaries (Hewson et al., 2001; Cissoko et al., 2008). Early studies in these systems have shown viral like particle (VLP) abundances and viral productivity (i.e., the number of viruses produced per hour) in rivers can be equivalent, or higher, than those in marine systems (Peduzzi and Luef, 2008; Corinaldesi et al., 2010; Rowe et al., 2012; Peduzzi, 2016). Additionally, early river studies found that up to 80% of bacterial isolate strains from sediments had virulent phage that could be isolated (Lammers, 1992). Together, these foundational works highlight the importance of viral predation in regulating microbial dynamics in river ecosystems.

There are two key reasons why it remains difficult to link viral communities to river ecosystem function. First, river microbiome studies are rarely genome-resolved, both from a bacterial and viral perspective. While there is still much to explore, most information on aquatic virus dynamics pertains to oceanic studies (Vincent and Vardi, 2023), and rivers are described as one of the most underexplored aquatic ecosystem with metagenomics, second only to glacier microbiomes (Chu et al., 2020). Although the taxonomic composition of microbial communities in rivers has been well-described by 16S rRNA gene amplicon surveys (Hou et al., 2017; Nelson et al., 2019), it remains unclear how microbial membership relates to relevant ecosystem processes. Likewise, our ability to link the viral community to their respective microbial hosts, and subsequently to ecosystem biogeochemistry, remains hindered by a lack of genome-resolved studies. Second, river studies are often temporally constrained. Although significant changes in river chemistry and hydrology are observed at seasonal periods (Tomalski et al., 2021), they are also known to change at sub-daily scales (Lundquist and Cayan, 2002; Alonso et al., 2017), particularly in human-impacted rivers affected by wastewater treatment plant effluent and reservoirs (Luo et al., 2020; Wang et al., 2021; Lu et al., 2022). Additionally, microbial communities double, evolve, and shift metabolically at an hourly basis (Wang et al., 2015; Erbilgin et al., 2017; Gibson et al., 2018). Nonetheless, river microbiome time-series are often resolved at seasonal scales (Kaevska et al., 2016; Malki et al., 2021), making our understanding of viral and microbial community dynamics across relevant short-scale gradients poorly understood.

To address these knowledge gaps, we collected a finely resolved metagenomic time-series at the River Erpe near Berlin, Germany, a lowland river receiving treated wastewater. Our sampling campaign included biogeochemical measurements every 3 hours for 48 hours across both surface water (SW) and pore water (PW) compartments that were paired to metagenomics and metabolomics **(Figure 1A-D)**. This study design provided a metagenomically resolved dataset which enabled us to interrogate how viral and microbial communities are structured across river compartments, and how this metabolic potential could modulate biogeochemical processes. Additionally, the unparalleled temporal resolution of our dataset allowed us to analyze both the persistence of viral and microbial communities across compartments, as well as the individual genome stability throughout the 48 hours of sampling. Finally, by using genome-resolved metagenomics, we show that viruses can be linked to hosts in river ecosystems, and that these linkages reveal putative interactions that may be central to ecosystem biogeochemistry.

**Figure 1:**
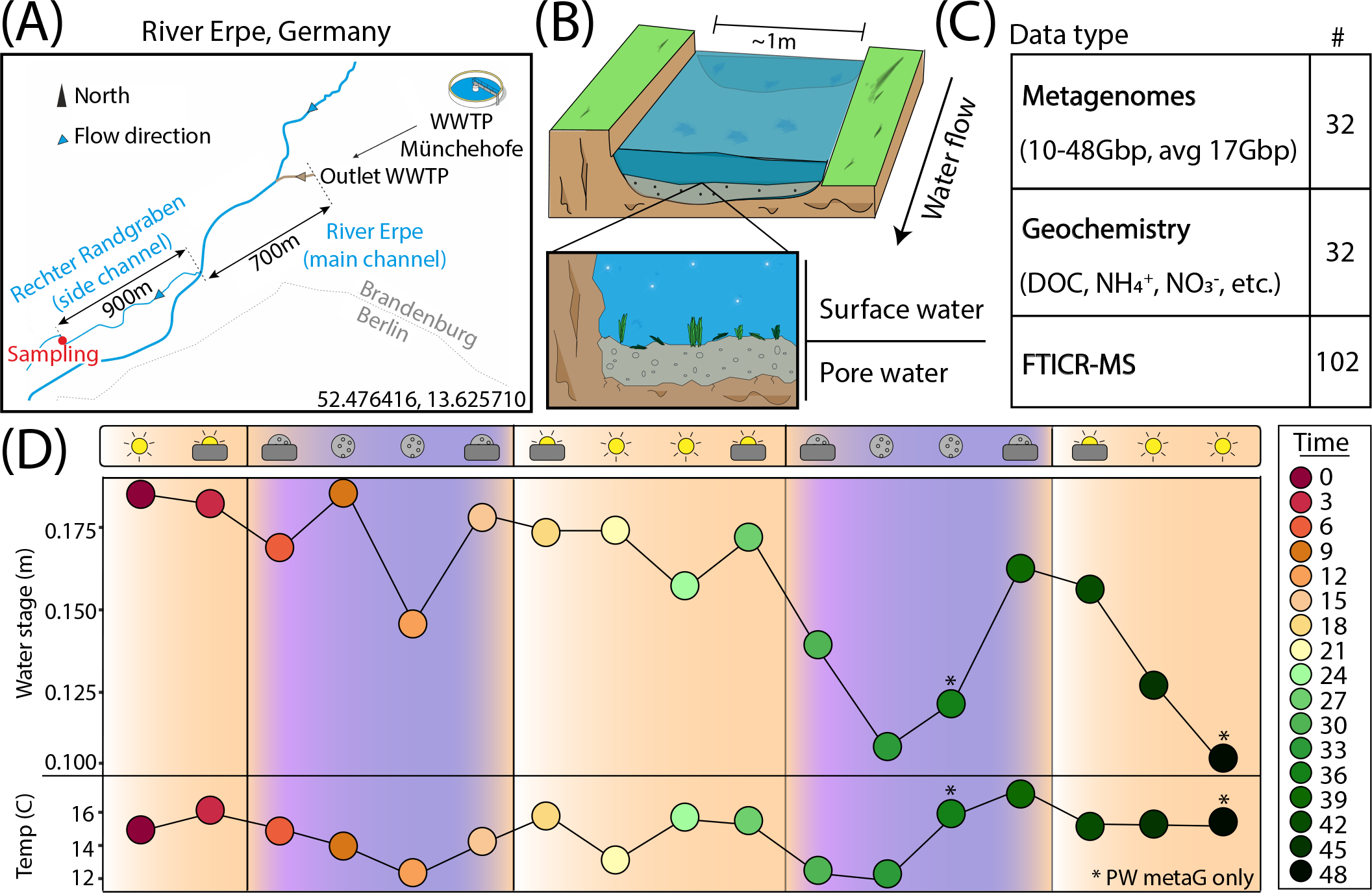
***Experimental design enables a genome- and time-resolved view of microbial communities at a finely scaled resolution.* A)** River Erpe sampling site that is located near Berlin, Germany. **B)** Conceptual schematic of the surface and pore water compartments that were sampled as part of this research. **C)** Table of data types that were collected as part of this sampling effort. **D)** Sampling schematic over 48-hour period with two ecological variables (water stage, and temperature) shown across the timepoints collected. The colors and icons highlight the hour of the day when samples were collected.

## Methods

### Sample collection, DNA isolation, and chemical characterization

The River Erpe is highly influenced by diurnally fluctuating effluent volumes of the Münchehofe wastewater treatment plant and consists of up to 80% treated wastewater (Mueller et al., 2021). Our sampling site is in a side channel with a mean discharge of 25 l/s (Lewandowski et al., 2011; Mueller et al., 2021) (**Figure 1A**). For sample collection, a sampling station was set up ∼1m from the shoreline of the River Erpe side channel “Rechter Randgraben” (52.476416, 13.625710), 1.6km from the wastewater treatment plant outlet leading the same water as in the main channel as previously described (Mueller et al., 2021), and in accordance to the Worldwide Hydrobiogeochemistry Observation Network for Dynamic River Systems (WHONDRS) protocol (Stegen and Goldman, 2018). Briefly, for surface water (SW), 60ml at a time of SW were collected manually with a syringe and tubing fixed in the water column and then passed through a 0.20μm filter until clogged. A cap was then put on the filter, filled with 3ml RNAlater, and refrigerated until extraction. For pore water (PW), 60ml of PW from 25cm sediment depth were collected with a stainless-steel rod in the middle of the channel. The rods were covered with a filter mesh sock over the screened area at the tip, pushed into the sediment, and equipped with a Teflon suction line. Samples were then taken by manually pulling 60ml of PW with syringes attached to the suction line and filtering them through a 0.20μm filter until clogged. The filter was then capped, filled with 3ml RNAlater, and refrigerated until extraction. Each of these processes were repeated every 3 hours over a period of 48hrs in September of 2018, resulting in 15 SW and 17 PW metagenomes. 2 SW samples failed due to lack of biomass. For DNA isolation, filters were cut into ∼5mm^2^ pieces and added to the bead bashing tubes of Quick-DNA Soil Microbe Microprep Kit (Zymo). The nucleic acids were then extracted according to the manufacturer protocol and sequenced at the Genomics Shared Resource Anschutz Medical Campus, Colorado. Accession numbers, total metagenomic reads, and sample sizes can be found on **Supplemental Table 1** and the original data repository (Wells et al., 2019).

Chemical characterization was performed as previously described (Mueller et al., 2021). Water samples were filtered with 0.2μm polyethersulfone Sterivex for Fourier transform ion cyclotron resonance mass spectrometer (FTICR-MS) analysis or regenerated cellulose for all other analytes, then acidified to a pH of 2 with 2M HCl and stored at -18°C until analysis. Samples were analyzed at the Leibniz Institute of Freshwater Ecology and Inland Fisheries for nitrate and sulfate (ion chromatography, Metrohm 930 Compact IC Flex), ammonium and soluble reactive phosphorous (SRP) (segmented flow analyzer Skalar SAN, Skalar Analytical B.V., Netherlands), and manganese and iron (inductively coupled plasma optical emission spectrometry (ICP-OES), (ICP iCAP 6000 series, Thermo Fisher Scientific Inc.). Dissolved organic carbon (DOC) concentrations were analyzed via infrared gas analyzer (NDIR) after combustion (TOC/TN Analyzer, Shimadzu). Dissolved organic matter (DOM) data is part of the WHONDRS dataset (Wells et al., 2019) and was analyzed using a 12T Bruker SolariX FTICR-MS (Bruker, SolariX, Billerica, MA, USA) at the Environmental Molecular Sciences Laboratory in Richland, WA. Once peaks were picked using the Bruker data analysis software and formulas were assigned using Formularity (Tolić et al., 2017), DOM was classified into seven compound classes based upon hydrogen to carbon ratio (H:C), and oxygen to carbon (O:C) ratios (Kim et al., 2003). FTICR-MS analysis does not allow for a quantitative approach, therefore compound class data was analyzed qualitatively, and DOM composition was evaluated using the number of molecular formulas in every compound class as described in the original publication (Mueller et al., 2021). The biogeochemical measurements for this study can all be found on **Supplemental Table 1**.

### Metagenome data processing and assembly

Each set of metagenomic reads were trimmed using Sickle v1.33 with default settings (Joshi NA, 2011), and assessed using FastQC (v0.11.2) (Andrews, n.d.). Trimmed reads were then assembled with either 1) metaSPAdes BBCMS pipeline (v3.13.0) (Metagenome Assembly Workflow (v1.0.1) — NMDC Workflows 0.2a documentation, n.d.), 2) Megahit (v1.2.9) (Li et al., 2015), or 3) IDBA UD (v.1.1.0) (Peng et al., 2012). For metaSPAdes pipeline, reads were merged into a single .fa file using fq2fa (Shen et al., 2016). Then, bbcms was run with flags “mincount = 2”, and “highcountfraction = 0.6”, followed by metaSPAdes using kmers 33, 55, 77, 99, 127, and flag “–meta”. For Megahit, reads were assembled with flags “k-min = 31”, “k-max = 121”, “k-step = 10”, and “m = 0.4”. For IDBA_UD, samples were rarefied to 25% of reads using BBMAP’s reformat.sh (Bushnell, 2014) with flags “samplerate = 0.25” and “sampleseed = 1234”. These 25% of subset reads were then merged into a single .fa file using fq2fa (Shen et al., 2016) and then assembled with default parameters. Assembly statistics for each sample can be found in **Supplemental Table 1**.

### Viral identification, taxonomy, and annotations

Viral metagenome assembled genomes (vMAGs) were identified from each set of assemblies using Virsorter2 and CheckV using the established protocols.io methods (Guo et al., 2021a, 2021b). Resulting genomes were then screened based on VirSorter2 and checkV output for viral and host gene counts, VirSorter2 viral scores, and hallmark gene counts (Guo et al., 2021b). Viruses were then annotated with DRAM-v using the “--use_uniref” flag, and further manually curated according to the established protocol (Shaffer et al., 2020; Guo et al., 2021b). The resulting subset of 6,500 viral genomes were clustered at 95% ANI across 85% of shortest contig per MIUViG standards (Roux et al., 2018) resulting in 1,230 viral populations.

Viral taxonomic identification of viral populations was performed using protein clustering methods with vContact2 using default methods (Bin Jang et al., 2019). We supplemented the standard RefSeq v211 database containing 4,533 vMAGs with viral genomes from an additional 303 river and wastewater treatment plant metagenomes that were publicly available from 1) JGI IMG/VR (6,254 vMAGs ≥10kb), 2) two previously unpublished anaerobic digestor metagenomic datasets that were mined in-house (14,436 vMAGs ≥10kb), 3) a previously published wastewater treatment plant sludge database (7,443 vMAGs ≥10kb) (Shi et al., 2022), 4) a previously available reference database that included freshwater ecosystem viruses (2,032 vMAGs ≥10kb) (Rodríguez-Ramos et al., 2022), and 5) the 43 TARA Oceans Virome datasets (5,476 vMAGs ≥10kb) (Brum et al., 2015). This resulted in an additional 35,641 reference vMAGs in our network. Proteins file for all vMAGs used in the network as well as accession numbers are available on Zenodo (*will be available upon publication*). Results from vContact2 can be found in **Supplemental Table 2.**

Viral population genome representatives were annotated using DRAM-v (Shaffer et al., 2020). To identify putative auxiliary metabolic genes (AMGs), auxiliary scores were assigned by DRAM-v to each annotated gene based on the following previously described ranking system: A gene is given an auxiliary score of 1 if there is at least one hallmark gene on both the left and right flanks, indicating the gene is likely viral. An auxiliary score of 2 is assigned when the gene has a viral hallmark gene on one flank and a viral-like gene on the other flank. An auxiliary score of 3 is assigned to genes that have a viral-like gene on both flanks (Shaffer et al., 2020; Rodríguez-Ramos et al., 2022). Genes identified by DRAM-v as being high-confidence possible AMGs (auxiliary scores 1-3) were subjected to protein modeling using Protein Homology / AnalogY Recognition Engine (PHYRE2) (Kelley et al., 2015), and manually verified. All files for vMAG quality and annotations can be found in **Supplemental Table 2.**

### Bacterial and archaeal metagenomic binning, quality control, annotation, and taxonomy

Bacterial and archaeal genomes were binned from each set of assemblies with MetaBAT v2.12.1 (Kang et al., 2019) as previously described (Rodríguez-Ramos et al., 2022). Briefly, reads were mapped to each respective assembly to get coverage information using BBmap (Bushnell, 2014), and then MetaBAT was run with default settings on each assembly after filtering for scaffolds ≥2,500bp. Quality for each MAG was then assessed using CheckM (v1.1.2) (Parks et al., 2015). To ensure that only quality MAGs were utilized for analyses, we discarded all MAGs that were not medium quality (MQ) to high quality (HQ) according to MIMAG standards (Bowers et al., 2017), resulting in 1,033 MAGs. These MAGs were dereplicated using dRep (Olm et al., 2017) at 95% identity, resulting in 125 MAGs. These 125 MQHQ MAGs were annotated using the DRAM pipeline (Shaffer et al., 2020) as previously described (Rodríguez-Ramos et al., 2022). For taxonomic analyses, MAGs were classified using the Genome Taxonomy Database (GTDB) Toolkit v1.5.0 on November 2021 using the r202 database (Chaumeil et al., 2019). Genome quality, annotations, and taxonomy are reported in **Supplemental Table 3**.

### Virus host linkages

To identify virus-host linkages, we used 1) CRASS (Direct Repeat / Spacer based) v1.0.1 (Skennerton et al., 2013), 2) VirHostMatcher (alignment-free oligonucleotide frequency based) v.1.0.0 (Ahlgren et al., 2017), and 3) PHIST (all-versus-all exact matches based) v.1.0.0 (Zielezinski et al., 2021). CRASS protocol and scripts used are described in detail on GitHub (see **Data availability**). VirHostMatcher was run with default settings, and the best possible hit for each virus was considered only if it had a d2* dissimilarity score of < 0.2. PHIST was run with flag “-k = 25”, and a PHIST hit was considered only if it had a significant adjusted p-value of < 0.05. To be classified as a virus-host linkage, a virus-host pair had to be predicted by the significant consensus of both VirHostMatcher and PHIST or a virus-host pair had to have a CRASS linkage. With this consensus method, CRASS links, which were always considered good hits, agreed across 60% of predictions at the Genus level, 80% of predictions at the Order level, and 87% at the Class level, suggesting high accuracy of consensus-only, non-CRASS linked virus-host pairs. All virus-host predictions are in **Supplemental Table 2**.

### Genome relative abundance and normalization

To estimate the relative abundance of each vMAG and MAG, metagenomic reads for each sample were mapped to a database of vMAGs or MAGs with Bowtie2 (Langmead and Salzberg, 2012) at an identity of 95%, with minimum contig coverage of 75% and minimum depth coverage of 3x. To normalize abundances for known temporal omics data biases (Coenen et al., 2020), we performed a library size normalization of abundance tables using TMM (Robinson and Oshlack, 2010). Given that PW and SW organism abundances were drastically different in magnitude, and that abundance zeroes across compartments are likely real zeroes, vMAGs and MAGs were considered to be present if detectable in at least 10% of samples in either compartment. Organisms detected in > 10% PW samples were labeled “pore”, organisms detected in > 10% SW samples were labeled “surface”, organisms > 10% PW and SW samples were labeled “both”, and organisms that were in < 10% SW and PW samples were removed. Based on these groups, the TMM abundances file was split into two different files, one for PW samples (n = 17) including “pore” and “both” organisms, and one for SW samples (n = 15) including “surface” and “both” organisms. Abundances for vMAGs and MAGs can be found in **Supplemental Table 2** and **Supplemental Table 3**, and specific commands can be found on GitHub.

### Temporal and statistical analyses

Temporal analyses were all performed in R with the TMM normalized abundances described above. To determine which environmental parameters were significantly driving differences across our compartments, we performed multiple regressions using envfit in the vegan R package (Oksanen et al., 2016) across multiple types of ordinations. Principal Coordinate Analysis (PCA) for biogeochemistry were done with vegan in R. Dissimilarities in community composition were calculated with the Bray-Curtis metric in vegan (Oksanen et al., 2016) for all vMAGs and MAGs that were present in >3 samples per each compartment. Nonmetric multidimensional scaling (NMDS) was then used with k = 2 dimensions for visualization. An analysis of similarity (ANOSIM) was performed using the base R stats package in order to determine community similarity between river compartments. PERMANOVA analyses were done in R using the adonis function from vegan. The NMDS ordinations of the vMAGs and MAGs were compared using the PROCRUSTES function in vegan. To visualize the relative contribution of each biogeochemical variable, we calculated the envfit vector using function ordiArrowMul and plotted them using ggplot. Shannon’s H’ were done using TMM normalized values with vegan in R. Species accumulation curves were done using the vegan function specaccum in R. All R code and files are available on GitHub.

To determine the relative stability of surface and pore water communities, we first calculated the differences in Bray-Curtis dissimilarity for each sample and its prior timepoint and then ran an unpaired t test to compare the mean differences across compartments with the vegan package in R. For assigning the persistence of the different genomes, we used previously established metrics to assess persistent (present in ≥ 75% of samples), intermittent (present > 25% <75% of samples), or ephemeral (present in ≤ 25% of samples) categories (Chow and Fuhrman, 2012). For establishing the abundance stability, we assessed the total number of samples in which each individual persistent genome fluctuated by ± 25% of the median relative abundance value across all samples. Then, using the established cutoffs by Fuhrman and Chow et al (Chow and Fuhrman, 2012)., we categorized our genomes as stable (shifting in ≤ 25% of samples), intermediately stable (shifting in > 25% < 75% of samples) and unstable (shifting in ≥ 75% of samples). Fishers exact test for count data was used for assessing the significance of difference in stability metrics using fisher.test from R base stats package. The enrichment analyses for AMGs were performed using a hypergeometric test between the total AMGs in our dataset and the individual groups of AMGs present in either compartment. The code used is available on GitHub. All temporal analyses and results are in **Supplemental Table 4**.

To reduce the complexity of our microbial data so we could link viral and microbial communities more concretely to ecosystem biogeochemical cycling, we applied a Weighted Gene Correlation Network Analysis (WGCNA) to identify which groups of organisms co-occurred using TMM normalized values in R with package WGCNA (Langfelder and Horvath, 2008; R Core Team, 2018). A signed hybrid network was performed with a combined dataset of MAGs and vMAGs on a per-compartment basis. For SW, we used a minimum power threshold of 14 and a minimum module size of 20. For PW, we used a minimum power threshold of 8 with and a minimum module size of 20. For both networks, a reassign threshold of 0, and a merge cut height of 0.3 were used.

To link the modules to ecosystem biogeochemistry, we performed sparse partial least square regressions (sPLS) on the groups of organisms in each module. sPLS were done using TMM normalized values of co-occurring communities that resulted from WGCNA above in R with package PLS (Chung et al., 2012). Subnetwork membership was related to the overall genome significance for nitrate as described in the WGCNA tutorials document (see GitHub code) using R and the WGCNA package (Langfelder and Horvath, 2008). Full code for WGCNA and SPLS are available on GitHub along with detailed instructions and input files. Visualizations for figures 6a, 7b, and 7c were made using RawGraphs (Mauri et al., 2017).

### Data availability

The datasets supporting the conclusions of this article are publicly available and collected as part of the Worldwide Hydrobiogeochemistry Observation Network for Dynamic River Systems (WHONDRS) collective sequencing project and are all publicly available on ESS-Dive (Wells et al., 2019). The 125 MAGs are deposited under (*will be available upon publication)* and in Zenodo (*will be available upon publication*). The raw annotations for each genome are deposited on Zenodo (*will be available upon publication*). The 1230 vMAGs have been deposited NCBI under the BioProject ID (*will be available upon publication)* and in Zenodo (*will be available upon publication*). Additionally, the dataset of freshwater and wastewater viruses we used to cluster to the HUM-V viruses is also hosted on Zenodo (*will be available upon publication*). Supplemental tables will be available upon publication or by request to corresponding author at the pre-print stage. All scripts, commands, and input data used for this manuscript are available at https://github.com/jrr-microbio/erpe_river.

## Results

### Metagenomics uncovers viral novelty and biogeography of River Erpe viruses

We sampled 17 pore water (PW) and 15 surface water (SW) metagenomes collected over a 48-hour period using a Eulerian sampling scheme (i.e., at a fixed location) and collected 565.5Gbp of paired metagenomic sequencing (10-48Gbp/sample, 17Gbp avg.) (**Figure 1, Supplemental Table 1**). Assembly of these samples revealed 6,861 viral metagenome assembled genomes (vMAGs), of which 6,500 vMAGs were ≥10kb in length and were subsequently clustered into 1,230 species-level vMAGs (**Supplemental Table 2**). The average vMAG genome fragment was 24,164bp (180,216bp max) in the PW, and 19,553bp (153,177bp max) in the SW (**Supplemental Table 2**). Viral MAG richness was consistently 8 times higher (p < 0.01) in the SW (845.0 ± 124.4) compared to the PW (108.3 ± 49.7) and likely drove differences (p < 0.01) in Shannon’s diversity (H’) recorded for the SW (SW = 6.05 ± 0.17, PW = 3.67 ± 0.49) (**Supplemental Figure 1**). In addition to our vMAGs, we identified 1033 metagenome assembled genomes (MAGs) that were dereplicated at 95% identity into 125 medium and high-quality genome representatives. Similarly, MAG richness was higher (p < 0.01) in the SW (SW = 62.6 ± 7.2, PW = 21.8 ± 9.0), and showed significantly different patterns (p < 0.01) in terms of Shannon’s (H’) (SW = 2.9 ± 0.17, PW = 2.6 ± 0.3) (**Supplemental Figure 1**). Together, these results highlight how metagenomic sequencing can be leveraged to successfully reconstruct representative viral and microbial communities across river compartments.

Viruses from freshwater systems are not well sampled in the databases commonly used for taxonomic assignment in viral studies (Elbehery and Deng, 2022). To determine the extent of novel viral diversity recovered, we mined additional set of 21,022 vMAGs from a variety of freshwater, wastewater, and marine samples and added this to the original vContact2 database (**Supplemental Table 2**, see Materials and Methods). We then performed protein clustering of our unique 1,230 viruses with this modified aquatic database, revealing 3,030 viral clusters (VCs). This network was composed of 19,623 nodes with 679,402 edges, which was simplified to only show protein clusters that contained at least 1 vMAG from this study (**Figure 2A-C**).

**Figure 2:**
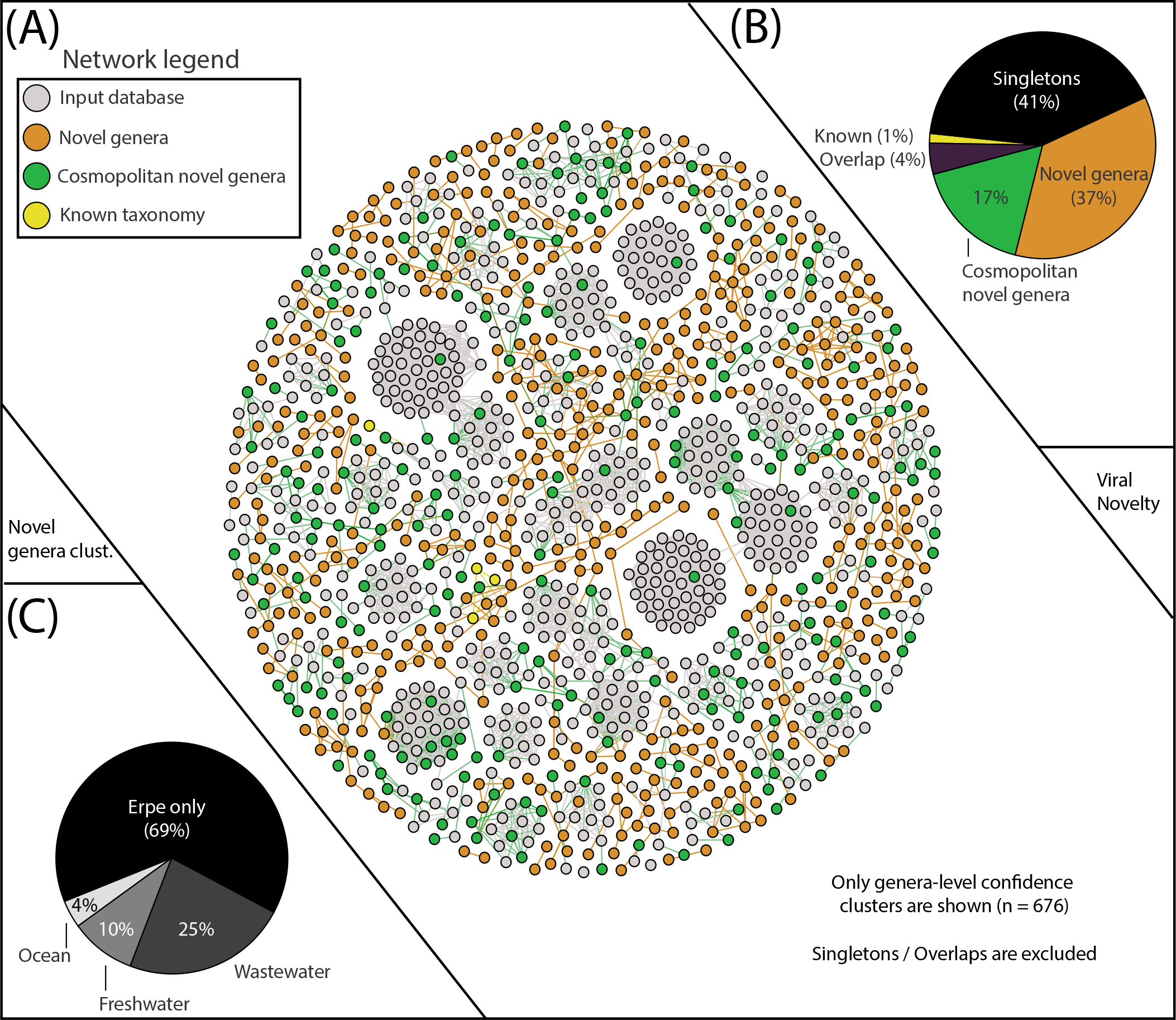
***vContact2 reveals Erpe vMAG database constitutes mostly novel genera, and a portion of these are cosmopolitan.* A)** vContact2 protein cluster (PC) similarity network where nodes represent vMAGs and edges show similarity across edges. Only high-confidence genera-level clusters are shown (n=676) with node color representing whether the vMAG pertains to our input databases (gray) or other categories assigned to vMAGs recovered here: orange shows novel genera (clustering only with Erpe genomes), green shows cosmopolitan novel genera (clustering with viruses from additional input database not from RefSeq), and yellow represents vMAGs with known taxonomy (clustering with known RefSeq vMAGs). Singletons (genomes that do not cluster with any other genomes) are excluded from the visualization (n=518). **B)** Pie chart shows the distribution of the different categories from the vContact2 network of vMAGs recovered. “Overlap” refers to a category where vContact2 assigns a vMAG to more than one cluster but cannot confidently place in either. **C)** Pie chart shows the proportion of vMAGs from novel genera in this study that were clustering with vMAGs from different environmental input databases.

Of our 1,230 vMAGs, 1% clustered to known taxonomic representatives of the Caudovirales Order (8 Podoviridae, 7 Siphoviridae, 3 Myoviridae). Of the remaining vMAGs, 37% clustered only to Erpe viruses, constituting 189 novel genera. An additional 41% did not cluster to any vMAG in our database and were “singletons” or “outliers”. Interestingly, 17% of our total vMAGs and nearly half of our novel genera were cosmopolitan in aquatic ecosystems, meaning that while these vMAGs failed to cluster with taxonomically known strains, they did cluster with vMAGs recovered from other ecosystems (**Figure 2B**). Specifically, our cosmopolitan novel genera clustered with vMAGs from wastewater treatment plant sludge or effluent (n=168), other rivers surface or sediment samples (n=65), and marine samples of the TARA oceans dataset (n=25) (**Figure 2C**).

### Viral and microbial river microbiomes are compartment-specific and coordinated with each other

The collected biogeochemistry was significantly structured across compartments and explained a large portion of the total variation in our samples (R^2^ = 0.79, p < 0.01) (**Figure 3A**). The surface water compartment was driven mostly by 1) the accumulation of alternative terminal electron acceptors (i.e., nitrate (NO_3_^-^), and sulfate (SO_4_^+^)), 2) the availability of nitrogen compounds (i.e., total nitrogen, avg. N), and 3) a more negative overall nominal oxidative state of carbon (NOSC) and a higher H:C ratio. Conversely, the pore water was characterized by 1) accumulation of NH_4_, 2) the availability of soluble reactive phosphorous (SRP), and 3) the overall concentration of carbon (avg. C), its aromaticity index (AI), and the quantity of double bond equivalents per molecule (DBE). In summary our redox data indicated more oxidative conditions in the SW while the FTICR-MS data showed that SW carbon was likely more labile, accessible, and thermodynamically favorable.

**Figure 3:**
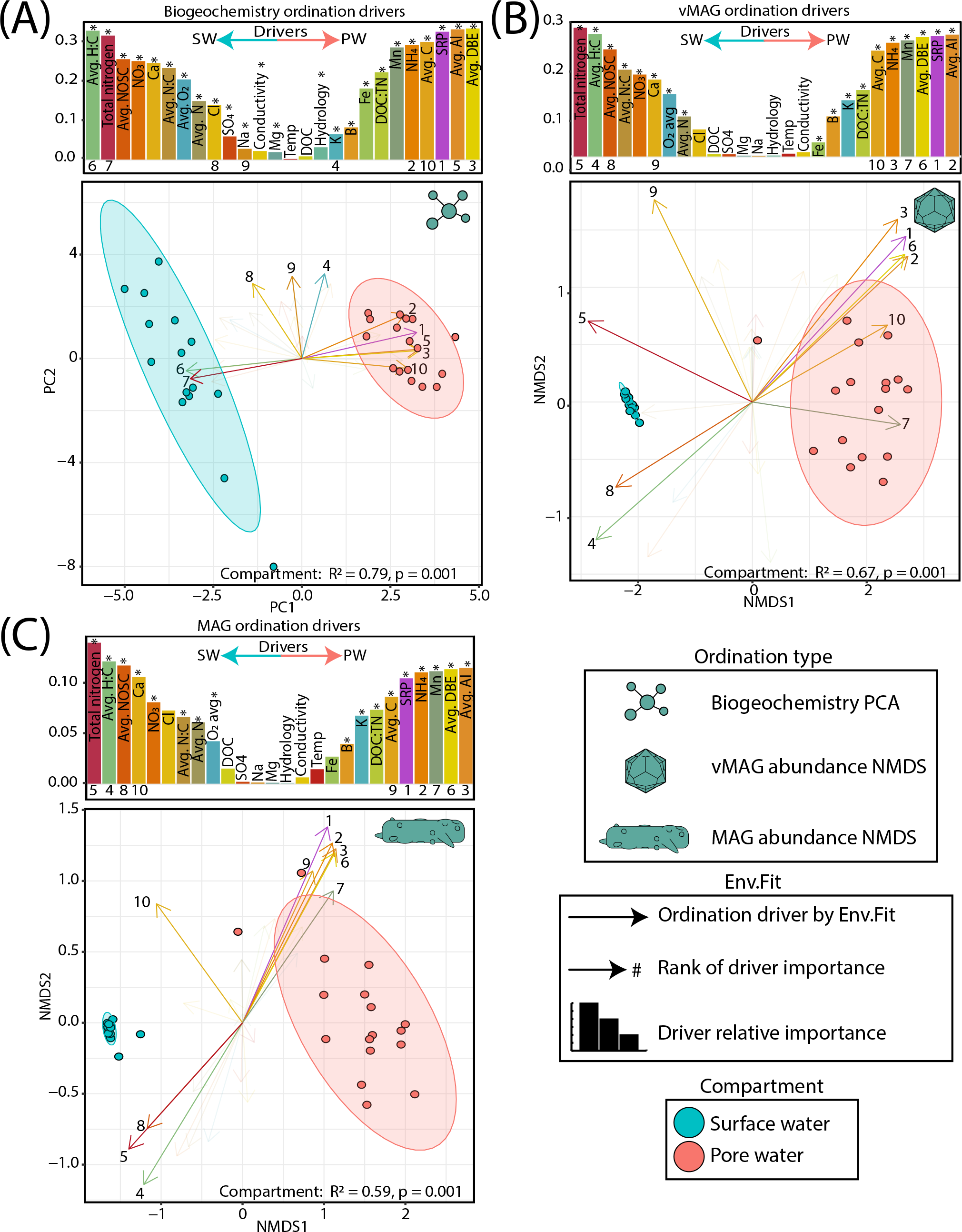
***Surface and pore water compartments have distinct viral communities and distributions are driven by biogeochemistry* A)** PCA plot of biogeochemical measurements where loadings and bars show the biogeochemical drivers per compartment. The size of bars represents the distance between the end of a loading arrow and the center of the plot. Within each bar plot, the drivers are labeled, and asterisks denote significant drivers by env.fit. The top 10 most significant drivers are numbered below each bar and are shown with solid, numbered arrows within the ordination below. **B)** NMDS ordination of river pore water and surface water vMAG abundances with bars and arrows showing the same as in (A). **C)** NMDS ordination of river pore water and surface water MAG abundances with bars and arrows showing the same as in (A). Non-compound abbreviations are: nominal oxidative state of carbon (NOSC), calcium (Ca), chlorine (Cl), sodium (Na), magnesium (Mg), dissolved organic carbon (DOC), soluble reactive phosphorous (SRP), aromaticity index (AI), and double bond equivalents (DBE). Note: NOSC values are plotted as the absolute value per value per sample (i.e., a higher SW NOSC driver value translates to a more negative NOSC measurement).

**Figure 4:**
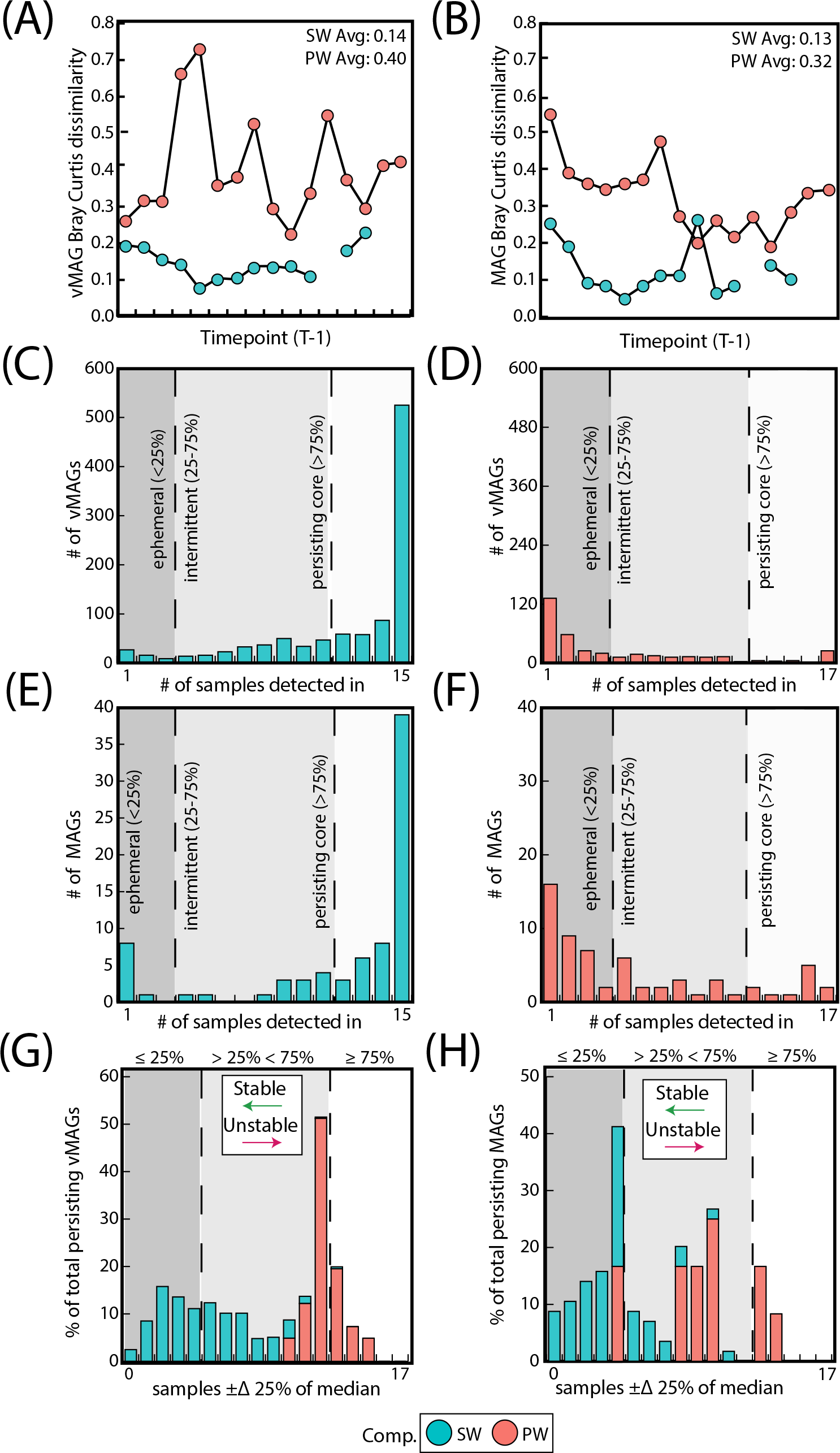
***Surface water communities are more stable and persistent than pore water communities.* A)** Difference in Bray-Curtis dissimilarities between each sample and its prior timepoint calculated for vMAGs and **B)** MAGs per compartment. **C)** Bar plots show the number of persistent, intermittent, and ephemeral vMAGs in the SW and **D)** the PW. **E)** Bar plots show the number of persistent, intermittent, and ephemeral MAGs in the SW and **F)** the PW. **G)** Bar plot where the x-axis shows the number of samples where each vMAG that fluctuates above or below 25% of their median values and the y-axis shows the normalized total percentage of persistent genomes per each compartment that are fluctuating. **H)** Identical bar plots to those in **G** but for MAGs.

To determine how viral and microbial communities were structured across these biogeochemical gradients, we recruited the time-series metagenomic reads to our viral database of 1,230 dereplicated vMAGs and 125 MAGs and then performed non-metric multidimensional scaling (NMDS) ordinations (**Figure 3B-C**). Like the geochemical PCA plots, PERMANOVA analyses showed that river compartment explained 67% (p < 0.01) and 59% (p < 0.01) of the variation in viral and microbial communities, respectively. The drivers of both viral and microbial communities were nearly identical in both magnitude and direction. Similarly, a PROCRUSTES analyses showed that vMAG and MAG ordinations are highly coordinated with each other (sum of squares = 0.027, corr. = 0.99, p < 0.01) emphasizing the dependencies between these communities (**Supplemental Figure 2**). Further highlighting these compartmental distinctions, the abundances of 85% of vMAGs (n = 1051) and 67% of MAGs (n = 87) were indicators of only one compartment (**Supplemental Table 5**). Interestingly, across both viral and microbial ordinations as well as our PCA, time only explained an additional 4-5% of the total variation, albeit significantly (p = 0.03, p = 0.02, and p < 0.01, respectively), likely due to long travel times and hydrological separation (**Supplemental Table 5**). Together, these results show viral and microbial communities are strongly structured by compartment, and that these in turn are structured due to compartment-dictated aqueous chemistry and the availability of suitable metabolic substrates.

### Temporally resolved metagenomics unveils compartment-level stability and persistence of viral and microbial communities

SW metagenomic temporal samples for both vMAGs and MAGs were on average 2-fold more similar than PW by Bray-Curtis dissimilarities (BC) (vMAG t = 6.3; MAG t = 6.2, p < 0.01) (**Figure 4AB**). We next evaluated whether the individual temporal persistence of the viral and microbial genomes shared similar patterns to the BC across compartments, and categorized members using persistence metrics that were previously established [84]. Briefly, if a viral genome was in more than 75% of the samples it was designated as persistent, between 25-75% of samples it was intermittent, and in less than 25% it was ephemeral. Of the 1,035 vMAGs detected in the SW compartment, 70% were categorized as “persistent”, with the remainder being 25% intermittent and 5% ephemeral. Contrastingly, of the 374 vMAGs detected in the PW, only 11% were categorized as persistent, with the remainder being 26% intermittent and 63% ephemeral (**Figure 4CD).** Similarly, the bacterial and archaeal MAGs shared comparable persistence patterns across the compartments (**Figure 4EF**). Combined, these results showed that SW communities were less temporally dynamic in terms of BC and had more persistently sampled genomes than the PW.

We then assessed whether the relative abundance of persistent genomes was also temporally stable. Based on Fuhrman and Chow (Chow and Fuhrman, 2012), we tallied the number of samples in which persistent vMAG and MAG relative abundances exceeded ± 25% of their respective median (**Figure 4GH, Supplementary Table 4**). Our results showed that both the relative abundance of vMAGs and MAGs in the SW fluctuate less over time than the PW as shown by Fishers exact t test (p < 0.01). Our persistence and temporal stability results supplement the observation that surface water communities in this urban stream change less over the 48-hour period than pore water communities which are more dynamic. Together, our results show that vMAGs and MAGs can be structured into temporally resolved groups at the genome level with varying stability.

### Genome-resolved virus-host analyses demonstrated viruses could infect highly abundant, phylogenetically diverse microbial genomes

We were able to predict hosts for 73 vMAGs, matching 30% of our total microbial genomes to a viral partner (**Figure 5)**. A majority (62%) of vMAGs with host associations were from the SW compartment, with 22% of host-associated vMAGs found in the PW, and around 10% found across both compartments. MAGs that had viruses linked to them were highly abundant, with 54% of our linked vMAGs infecting hosts of the top 25% most abundant MAGs. At the phylum level, 11 of the 20 identified phyla had evidence for a viral host. Notably, all the phyla that could not be assigned a viral link had 2 or less MAG representatives, with the exception of *Desulfobacterota* which had 6 MAGs. Additionally, of the 51 *Patescibacteria* MAGs we recovered in this study, we uncovered 12 possible viral genome links, which to our knowledge is one of the few reports of possible infective agents for members of this phylum (Holmfeldt et al., 2021; Trubl et al., 2021), and is the only one thus far reported in rivers. Ultimately, nearly a third of the genera from our MAG database as defined by GTDB were successfully linked to a vMAG, providing further evidence that viral predation is likely pervasive across these river microbial communities.

**Figure 5:**
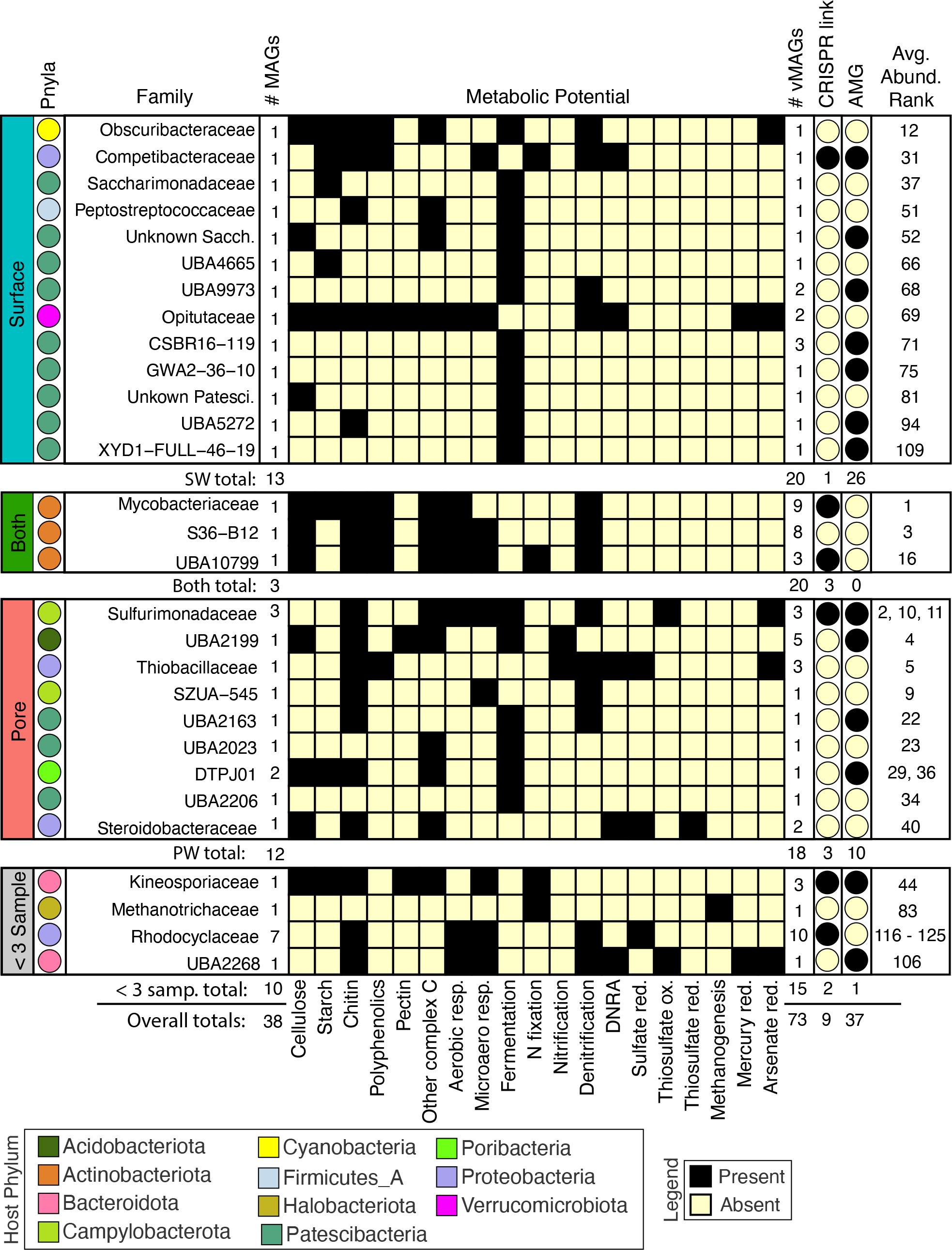
*Viruses infect abundant microorganisms in rivers which can influence aerobic and anaerobic C, N, and S cycling by predation or auxiliary metabolic genes.* MAG families that had a linkage to a virus are shown and split into their compartment-level distributions. From left to right: Colors of each circle on the leftmost side represent the Phyla, and for each family the total number of MAGs are shown. The presence absence heatmap describes the metabolisms of each family. Following the heatmap are the number of vMAGs that are linked in each family, whether the virus-host link is predicted by CRISPR or consensus method, and if at least 1 infecting vMAG with an AMG is reported. Numbers below each bounding box show totals of above criteria. The overall average rank of each MAG within a family is shown in the rightmost column.

To decipher the potential impacts that viral predation could have on ecosystem biogeochemical cycling, we metabolically characterized the 38 viral-linked MAGs from our genome-resolved database and saw a wide array of metabolisms spanning ecosystem redox gradients (**Figure 5**). Across both compartments, viruses were inferred to impact hosts that could modulate both aerobic and microaerophilic metabolism (carbon respiration), as well as anaerobic metabolisms (nitrate reduction, fumarate reduction, fermentation, and nitrogen fixation). For example, vMAGs were predicted to infect hosts with metabolisms such as methanogenesis (e.g., *Methanothrix*), and sulfur metabolisms (e.g., *Sulfurimonas*), which were encoded more predominantly by MAGs in the PW. Interestingly, 20% of the vMAGs that infected hosts were cosmopolitan with representatives identified in other freshwater and wastewater systems. Together, these results suggest that viruses can infect hosts that can play key roles in ecosystem biogeochemical cycling, and insinuate that these influences are likely widespread across a wide biogeographical range.

### Virally encoded auxiliary metabolic genes can potentially alter host metabolic machinery

In addition to the impact on microbial communities via predation, viruses can also mediate biogeochemical cycles through enhancing host metabolism with Auxiliary Metabolic Genes (AMGs). We mined our 1,230 vMAGs for putative AMGs and found 165 unique viral AMG candidates after quality filtering. We failed to see a statistical enrichment for the number of AMGs in either compartment (Fisher’s exact p = 0.77), indicating their shared importance. The functionalities of these AMGs at the gene annotation level (e.g., KO number) were mostly conserved across compartments, with only 27% of unique gene IDs present in both compartments. However, at the functional module level (e.g., amino acid metabolism) 69% of metabolisms were present across both our ecological gradients (**Figure 6B**). Conserved DRAM categories across compartments pertained to electron acceptor utilization (e.g., oxygen, nitrate), carbon utilization (e.g., CAZyme inferred substrates), and other reactions (heavy metal usage, nitrification). We note that genes necessary for viral replication like nucleotide biosynthesis, ribosomal proteins, host mimicry, glycan biosynthesis, cofactor and vitamin metabolism, and molecular transporter were conserved between compartments.

Nonetheless, there were some AMGs that did show compartment specificity. Within the surface water, we detected AMGs involved in organic nitrogen mineralization and transcriptional regulation (e.g., peptidases (M50)), sugar metabolism (e.g., fructose, mannose), and motility (e.g., flagellar assembly). These unique AMGs could be associated with a lifestyle supported by more favorable carbon, more aquatic environments that would favor mobility, and differences in protein content associated with the SW environment. On the other hand, in the pore water we detected AMGs that encoded for plant hemicellulose degradation and cobalamin biosynthesis, adaptations that could sustain metabolism in a litter impacted, anoxic habitat. Ultimately, these results suggest that like their microbial hosts, viral AMGs can potentially have some degree of functional tuning or filtering by environmental conditions.

We next considered AMGs that either expanded the host metabolism or that were complementary to the host metabolism (i.e., Class I AMGs) (Hurwitz and U’Ren, 2016). Of the 12 *Patescibacteria* MAGs that had possible viral genome links, MAG representative CSBR16-119 had two possible vMAG linkages. A comparison of the metabolic capabilities of the host and viral genomes indicated multiple shared genes (**Figure 6C, Supplemental Table 2**). For example, a peptidase-like protein (M50) that is inferred transcriptional regulator (Rawlings et al., 2018) was present in both the Patescibacteria MAG and its infecting vMAG and shared 77% nucleotide and 99% amino acid similarity across the length of the open reading frame (**Supplemental Table 2**). The microbial host genome also had a single copy of ribosome L28 encoded, and two viral genomes putatively infecting it that contained 1 AMG each for the ribosomal protein L28. Both L28 AMGs shared 93% nucleotide identity (90%, and 100% query coverage) (**Supplemental Table 2**). It is possible that these viral proteins show strong homology to replace the cellular versions in the host (Mizuno et al., 2019), or can function to benefit the host, potentially leading to growth rate enhancement (Brahim Belhaouari et al., 2022).

A second putatively infected persistent genome was *Proteobacteria* UBA2383 (a novel unclassified *Competibacteraceae*) which had broad metabolic capabilities (**Figure 6D**). This MAG was inferred to be a facultative aerobe encoding genes for aerobic respiration and for denitrification. The genome encoded genes supporting a heterotrophic lifestyle including CAZymes necessary for the degradation of complex carbon substrates like chitin, starch, and polyphenol, and complete glycolytic and TCA pathways for oxidation of carbon. This MAG also encoded the ability to fix nitrogen. The two vMAGs that were associated with this genome encoded genes to support this latter metabolism, e.g., GTP cyclohydrolase, which generates important co-factors for the nitrogen fixation process (**Supplemental Table 2**) (He and Rosazza, 2003). Additional AMGs encoded by infecting viruses could potentially enhance nucleotide biosynthesis (dCTP deaminase, dUTP pyrophosphatase, thymidylate synthase) as well as other viral functions like host mimicry genes (i.e., 7-cyano-7-deazaguanine synthase, 6-carboxy-5,6,7,8-tetrahydropterin synthase) to avoid the CRISPR defense mechanisms encoded within the host *Proteobacteria*. Together, these results show that viruses are not only important due to viral predation in river ecosystems, but that they can also potentially play critical roles in reprogramming host metabolism.

### Co-occurrence networks elucidate ecological patterns that inform ecosystem biogeochemistry

To link viral and microbial communities more concretely to ecosystem biogeochemical cycling, we applied a Weighted Gene Correlation Network Analysis (WGCNA) to identify which groups of organisms co-occurred over the 48-hour sampling time. Highlighting the clear distinctions in SW and PW compartments, WGCNA analyses could not be reasonably performed simultaneously on a dataset containing both SW and PW communities (scale free topology model fit max = 0.32 at power = 20). As such, using only microbial and viral genomic abundances from either SW or PW separately, we identified 15 and 4 co-occurring modules in the SW and PW, respectively (**Supplemental Figure 4**). The largest module in both networks (turquoise module) contained 254 genomes in the SW and 71 in the PW. In the SW compartment, the overall modules had an average richness of 66 vMAGs and 5 MAGs, while in the PW they had an average richness of 46 vMAGs and 10 MAGs.

Overall, both surface and pore water communities had modules of co-occurring genomes that were significantly related (R^2^ > 3, p < 0.05) by sparse partial least square regressions (sPLS) to the collected biogeochemical measurements (**Figure 7A**). Only total Fe concentrations were related to modules in both the SW (brown, salmon modules) and PW (red module). SW modules were uniquely related to variables pertinent to nitrogen (nitrate, average total nitrogen), carbon (average total carbon, aromaticity index, hydrogen:carbon), as well as physical (temperature, water stage) and geochemical (magnesium, calcium, manganese, ammonium, sulfate) features in these samples. Of the 8 modules that were significantly related to ecosystem hydrobiogeochemical features, viruses had significant variable importance in projection scores (VIP > 1) in 7 of them, and 70% of the most significantly related genomes across all regressions were viral (**Figure 7B**).

**Figure 6:**
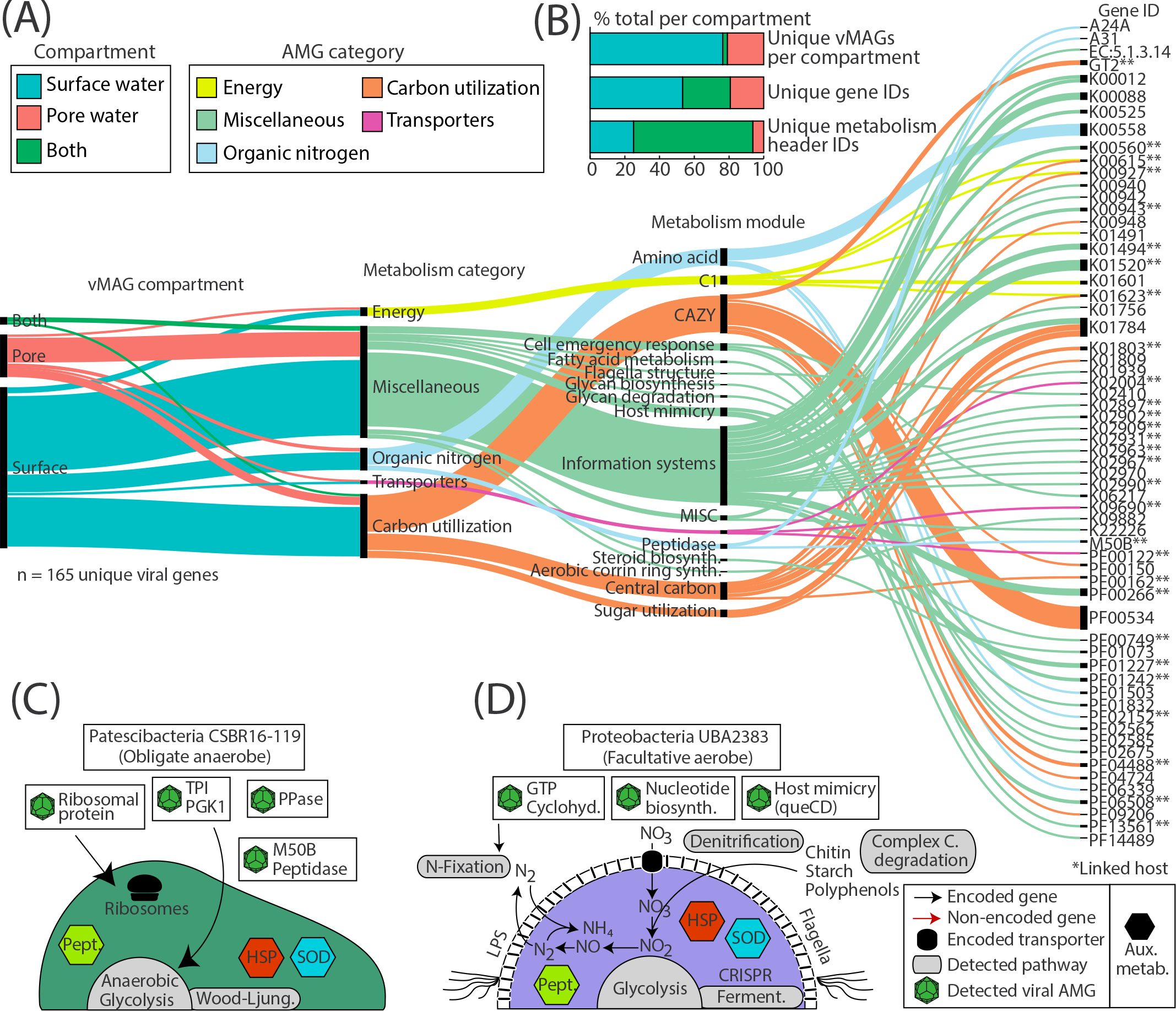
Distribution of viral Auxiliary Metabolic Genes (AMGs) and their function reveals key viral interactions that can enhance host metabolism in river ecosystems. A) Alluvial plot shows the subset of AMGs (77%, n=165) that had a metabolic function annotated by DRAM-v and were 1) not at the end of a contig and 2) did not contain a transposon like element. In the first vertical line, colors show the compartments that each vMAG with an AMG was detected in. The second vertical line shows the different DRAM-v metabolic categories for each AMG. The next vertical line shows the specific metabolic module name as categorized by DRAM. The final line contains each of the Gene IDs for the detected AMGs. Genes that can have multiple functions (n = 13) are duplicated and treated as individual genes within each category. **B)** Stacked bar charts show the proportion of total AMGs encoded in vMAGs from different compartments at the scaffold, gene ID, and metabolism header ID level as shown in **(A)**. **C-D)** Genome cartoons of two computationally linked bacterial hosts and their respective metabolisms. Detected viral AMGs are shown as viral icons above each genome cartoon. Pept. = peptidases, HSP = heat shock proteins, SOD = superoxide dismutase, queCD = 7-cyano-7-deazaguanine synthase, 6-carboxy-5,6,7,8-tetrahydropterin synthase.

**Figure 7:**
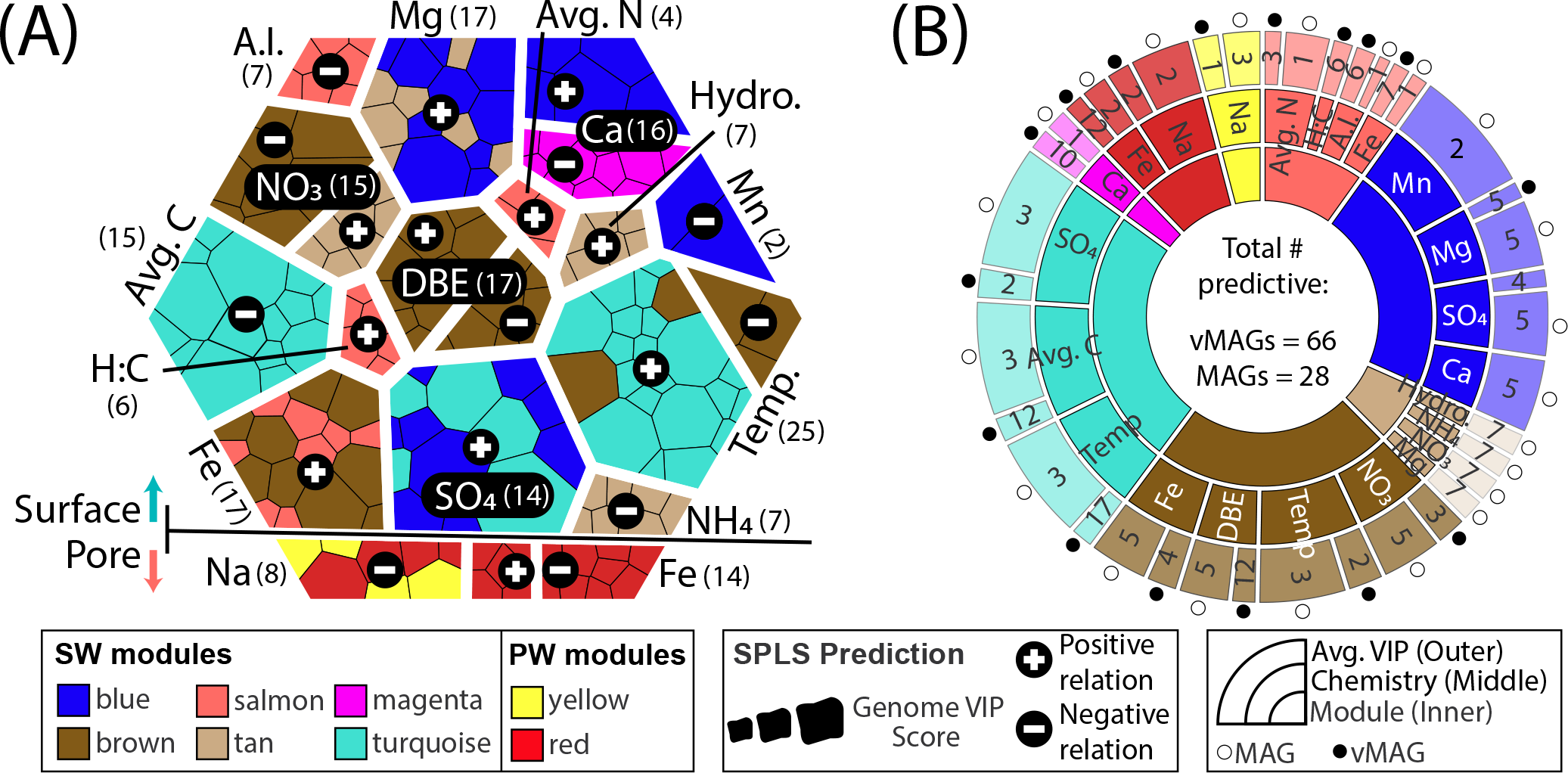
**WGCNA co-occurrence networks reveal ecologically similar groups that are related to overall ecosystem biogeochemistry**. **A)** Voronoi diagram shows VIP values of predictions for each predictive genome using a hierarchy structure. Each amorphous square within a group represents a single MAG or vMAG. At the first level (i.e., splitting of the large hexagon into upper and lower groups), SW (top) and PW (bottom) predictions are shown. At the second level (i.e., grouping of individual chemical variables predicted across each compartment), individual chemical variables are shown, per each compartment, and how many vMAGs/MAGs were predictive are denoted by numbers next to each variable name. At the third level (i.e., individual amorphous square or genomes), shapes are sized by the VIP score (>1) of genomes that predict that variable and are colored by their respective WGCNA module. **B)** Sunburst diagram shows the predictive WGCNA modules in the innermost level, followed by what chemical values each module predicts in the middle level. The outer level shows the average variable importance in projection (VIP) score for each genome type: vMAG (black circles) and MAGs (white circles) for that chemical prediction.

Of the 73 vMAGs and 38 MAGs that were computationally linked (**Figure 5, Figure 6**), nearly a quarter of those vMAGs and a third of MAGs were grouped into the same co-occurring modules. Interestingly, the SW brown module was related to the total nitrate concentrations in our dataset and contained a co-occurring virus-host link (**Figure 8A**). The host genome was the *Competibacteraceae* genome in **Figure 6D** and its putatively infecting a virus, which together could play roles in modulating the nitrogen cycling through both fixation and denitrification. This virus and microbial host pair had significant negative correlations to nitrate concentrations and were the second and fourth most significantly related genomes to nitrate within the brown module. The virus bacterial ratio (VBR) for these two organisms was nearly 1:1 and significantly correlated, which is expected of kill the winner dynamics (Trubl et al., 2021), and ultimately highlighting the possible dependency of an infecting vMAG and its host (**Figure 8B**). In support of this relationship, the viral genome coverages were on average 10x more than the putative host MAG coverage, suggesting a possible lytic infection lifestyle. Further underlining the importance of these related genomes, both were designated as persistent (i.e., present in >75% of all collected timepoints) and were the 1^st^ (vMAG) and 9^th^ (MAG) most abundant genomes detected in the surface waters. Together, these results indicate that viral and microbial predation dynamics are strongly related to, and could potentially influence, ecosystem biogeochemistry which suggests viral content may be useful units for modeling river ecosystem biogeochemistry.

**Figure 8:**
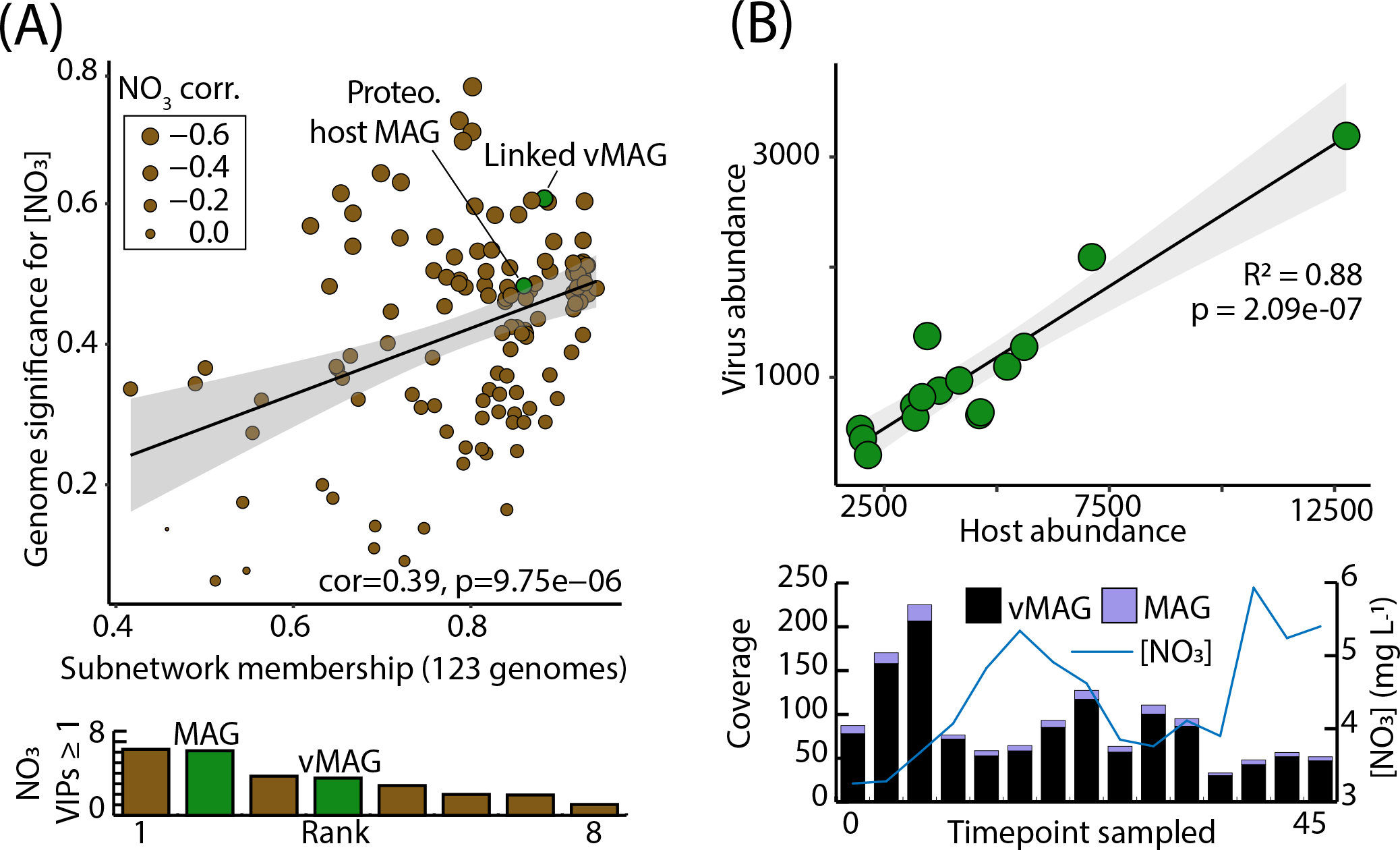
Computationally linked vMAG and MAG pair that share co-occurrence patterns demonstrate high significance for nitrate, and display kill-the-winner dynamics. A) Scatterplot depicts the genomic significance for nitrate of each of the genomes in the brown module in relation to the membership of those genomes within the WGCNA network modules. Below, bar charts show the VIP score (≥1) of the different organisms in the brown module. **B)** A Virus bacteria ratio (VBR) plot of a viral genome within the brown module that was predicted to infect a Proteobacteria genome. Below it, bar plots show the total coverage across all samples for both the vMAG and the MAG, and a line graph shows the measured nitrate concentrations that these genomes predict.

## Discussion

### Viral reference databases underrepresent certain habitats, missing cosmopolitan, ecologically relevant lineages

Within the 48-hour time-series metagenomes, we identified 1,230 dereplicated vMAGs that spanned surface water (SW) and pore water (PW) compartments. The large majority of vMAGs from our dataset were of unknown taxonomy (99%), a finding previously reported in another genome resolved viral focused publication from river sediments (Rodríguez-Ramos et al., 2022). Together these two studies surmise the current state of genome-resolved viral analyses from river systems, highlighting how underexplored river ecosystems are from a viral perspective (Chu et al., 2020). Given that these recent viral river genomic studies have yet to be incorporated into viral reference datasets, as well as the influence of wastewater treatment on this urban stream, we tested whether adding more viral representatives (n = 21,022) from relevant ecosystems (i.e., wastewater treatment plant effluent, freshwater viruses, and ocean viruses) to our analyses could expand the relevance of the viruses recovered here (**Supplemental Figure 3**). Adding these additional genomes reduced the total number of River Erpe vMAGs that were categorized as singletons or outliers, resulting in the addition of 49 novel genera, and giving biogeographical context to 164 novel viral genera (**Figure 2A**).

The biogeography of viruses and how they are structured across spatial gradients has been studied previously in oceanic ecosystems (Roux et al., 2016; Jian et al., 2021), developing the idea of “global” or “core” viruses (Ignacio-Espinoza and Sullivan, 2012; Hurwitz et al., 2015; Kieft et al., 2020; Heyerhoff et al., 2022). Further, it has been shown that factors like the presence of AMGs can enable a virus to potentially survive across a variety of different ecosystems (Sullivan et al., 2006; Chow and Suttle, 2015). Nearly a quarter of our Erpe viruses formed genus-level clusters with viruses from wastewater and freshwater systems, and of those, 11% encoded a putative AMG with functions for metabolisms such as carbon utilization, organic nitrogen transformations, and housekeeping functions (i.e., transporters and flagellar assembly). While the protein clustering of River Erpe vMAGs to wastewater viruses was not entirely surprising given the sampling location was downstream from the wastewater outlet (Mueller et al., 2021), we note that we also clustered a similar proportion of viruses to other viral genomes from river systems. Notably, this similar clustering proportion for River Erpe viruses was not observed with the TARA ocean viruses (**Figure 2C**, **Supplemental Figure 3**). These results hint at possible ecosystem filtering that may affect the biogeographical patterns of freshwater viruses. Our results also underscore the importance of customized, ecosystem relevant databases in environmental viromics for extending the ecological relevance of these ecosystem modulators, and further understanding the major drivers for river microbiomes.

### Temporally and spatially resolved metagenomics demonstrates that viral and microbial communities are compartment specific, and more stable in surface water than sediment pore water

Rivers are characterized by containing distinct ecological compartments that are in relatively close proximity to each other (e.g., surface waters and pore waters) (Stegen et al., 2018). Furthermore, river microbiomes have been reported to have temporal dynamics that can change at hourly or sub-daily scales (Lundquist and Cayan, 2002; Alonso et al., 2017; Tomalski et al., 2021). Nonetheless, most river microbiome studies focus on a single compartment, and/or are examined at seasonal or yearly scales thus lacking relevant temporal resolution. To address this temporal knowledge gap, we sampled both the SW and PW compartments over a 48-hour period, with sampling every three hours. Within our identified vMAGs and MAGs, we saw very clear structuring of communities at this compartment scale and observed differences in their persistence and stability (**Figure 3ABC, Figure 4**). Like the sampled biogeochemistry, microbial and viral communities were heavily structured by river compartment (67% and 59%, respectively). These data support previous studies within urban rivers (using 16S rRNA gene sequencing) that have shown that chemistry like phosphorous, nitrate, and metals covaries with microbial communities across compartments (Wang et al., 2018).

Based on the inferred redox and our metagenomic data, we infer that the SW compartment microorganisms are preferentially utilizing oxygen, likely contributing to the accumulation of alternative terminal electron acceptors within this compartment (i.e., nitrate (NO_3_^-^), and sulfate (SO_4_^2-^)). Likewise, genomes with the capacity for anoxic metabolisms like methanogenesis, as well as sulfur reduction, were only detected in the porewater. In addition to redox features, our FTICR-MS data was also shown to be an important driver of both microbial and viral communities (**Figure 3**). Most of our FTICR-MS metrics indicated more labile and accessible carbon in the SW (lower H:C ratio, lower C:N ratio, lower DBE, lower AI) (Seo et al., 2009; Bae et al., 2011; Ghosh and Leff, 2013; Mentges et al., 2017), however the NOSC values showed the opposite, with a more negative value being associated with the SW (Boye et al., 2017). Explaining these unexpected differences, it has been recently shown that variables other than NOSC may carry higher importance for predicting carbon lability (Garayburu-Caruso et al., 2020), particularly in environments with oxic conditions (Pracht et al., 2018). Ultimately these findings highlight the need for larger, cross river studies to decode the unifying factors, like redox and carbon quality, that control the structure and function of river microbiomes accounting for variables like river order, geography, time, and compartment.

Sampling with a Eulerian method allowed us to detect microbiomes passing through the same space over time in the SW and PW samples. Due to the flow rate of SW, and the potential that PW communities may be more biofilm impacted, we might have expected to see greater microbial and viral changes in the surface compartment than the sediments over the sampled time period. On the contrary, both vMAGs and MAGs were more persistent and had more stable abundance patterns over time in the SW of the River Erpe (**Figure 4AB**). One possible explanation could be methodological due to the PW being sampled less completely due to genomic extraction bias caused by fine grain sediments, less sampling volume, or strain level complexity. However, our species area curves did not signify an obvious difference in sampling exhaustion between these compartments (**Supplemental Figure 1**), leaving open the possibility that this finding may be biological. A possible biological explanation is that the strong influence of the wastewater treatment plant, where inputs were relatively uniform and continuous over time (Mueller et al., 2021), could contribute to the increased temporal stability we observed. It is also possible that the mixing in the PW hyporheic zone was more frequent that the flow rate within this channel. In support of the former, we did observe strong clustering between our viral genomes and wastewater treatment viral genomes (**Figure 2**). Importantly, our study validates other research indicating that surface water microbiomes are not unstable, or intractable (Graham et al., 2017), and could thus be important for the poorly resolved indices of river health and biogeochemistry that currently exist.

Ultimately, given the limited number of genome-resolved metagenomic studies in rivers, our results demonstrate that increasing the number of temporally resolved, multi-omic datasets could lead to the development of theoretical frameworks for understanding river viral and microbial community dynamics in natural and urban impacted rivers. Further, our results suggest that studies performed at these finely resolved scales can be informative of viral and microbial metabolisms, which are known to be critical for overall river respiration (Boulton et al., 1998; Newcomer et al., 2018). Finally, our results support the idea that understanding these dynamics is useful for enacting efficient sampling campaigns that can maximize efforts based on relevant biological and biogeochemical factors (i.e., adjusting frequency and depth of metagenomic sampling based on empirical knowledge of temporal microbiome dynamics, where perhaps SW communities require less efforts than PW communities).

### Viruses have the potential to regulate river biogeochemical cycles by predation and metabolic reprograming of microbial hosts

Although river viral ecology is only recently becoming appreciated, early works have suggested that viruses likely play key roles in the structuring of river microbial communities (Peduzzi and Luef, 2009; Peduzzi, 2016). By using a combination of computational methods, we were able to link 73 vMAGs to 38 MAGs spanning a wide range of taxonomic identities (**Figure 5**). One of the PW vMAGs in our dataset was identified to putatively infect an archaeal genome of the genera *Methanothrix*, a known canonical methanogen that accounts for the majority of *mcrA* transcripts in other freshwater environments (Angle et al., 2017). In addition to virally impacted microbial methane metabolisms, we show 11 of the 29 microbial families that were linked to a virus had the metabolic potential for denitrification, which could have ramifications for nitrous oxide emissions (Hu et al., 2016) (**Figure 5**). Together, our genome-resolved database of microbial metabolisms, and their putatively infecting viruses gives insight into the underpinnings of river metabolisms, including climate-critical metabolisms (i.e., nitrous oxide and methane production), and show that river microbiome studies can provide critical perspectives for understanding the impact that viruses can have in river ecosystems.

In addition to predation, viral auxiliary metabolic genes are recognized across aquatic systems to play key roles in host metabolic reprogramming and can encompass a wide range of processes from photosynthesis to the oxidation of sulfur (Sullivan et al., 2006; Anantharaman et al., 2014; Rodríguez-Ramos et al., 2022). We add to the existing literature and show AMGs in urban river systems may also impact redox important reactions involving nitrogen (peptidases), carbon (CAZymes), and sulfur (thiosulfate reduction) (**Figure 6**). Additionally, one of the vMAGs that was predicted to infect a *Patescibacteria* genome encoded a ribosomal protein that was similar to the gene encoded in the host genome (**Figure 6C**). Candidate phyla radiation (CPR) organisms like *Patescibacteria* are well known to be present in wastewater treatment plants (Wang et al., 2023), an idea supported by the fact 90% of the *Patescibacteria* genomes in our datasets originated from SW metagenomes. Interestingly, CPR organisms are also known to contain non-redundant, highly streamlined small genomes (Tian et al., 2020; Wang et al., 2023). As such, our results hint at the possibility that *Patescibacteria* viruses could help maintain those small genome sizes by encoding for necessary host genes, a concept with ecological precedence that has been previously demonstrated for the virus-host dependency of cyanobacterial photosynthesis in oceans (Sullivan et al., 2006).

While there are extremely limited studies from rivers examining viral AMGs, a prior study showed that river viruses contain AMGs that are expressed and potentially influence river biogeochemical cycles (Rodríguez-Ramos et al., 2022). Other works looking at vMAGs from freshwater lakes and estuaries have shown that some viruses seem to exhibit endemism for certain environments, meaning their distribution is limited to a small geographic area (Ruiz-Perez et al., 2019). This points to an interesting idea that perhaps AMGs are also tuned to the specific ecological functions of the sampled habitat, and as such that we could expect some degree of endemism. Interestingly, although individual AMG genes in the River Erpe were unique across compartments (27%), their functional categories were highly similar (69%) (**Figure 6B**). Our finding is different to a study recently reported form an estuary, where significant partitioning of AMG functions was reported between habitat types (water particle and sediment) (Luo et al., 2022). It is possible that due to the constant mixing of surface and HZ water in rivers, stratification at the genomic potential may be less notable, and expression information may be necessary to capture habitat specific differences. Lastly, this study highlights how moving forward annotation resolution and expanding reference database(s) are important factors to consider when extrapolating AMG inferences across datasets (Hurwitz and U’Ren, 2016; Shaffer et al., 2020).

By combining our computational virus-host linking and our AMG analyses with weighted correlation network analysis (WGCNA), we were able to link a specific virus-host pair to their potential impacts on nitrate utilization (**Figures 7 and 8**). A virus linked *Competibacteraceae* genome was detected to co-occur within the brown module with its infecting virus, and this module was highly negatively related to nitrate concentrations. This bacterial genome had the metabolic capability to fix nitrogen as well as denitrify (**Figure 6D**), and the latter metabolism could explain the negative relationship between this bacterium (and its crispr-linked virus) to SW nitrate concentration. In addition, we also provide AMG evidence that this virus could further alter host cell nitrogen use through modulating nitrogen fixation. Interactions like these are not unheard of in natural systems (Hurwitz and U’Ren, 2016), and ultimately suggest that viruses in river systems are possibly top-down (by predation) and bottom-up (by resource-control) regulators of ecosystem biogeochemical cycles.

In conclusion, how microbial and viral communities interact and change across spatial, and finely tuned temporal gradients is poorly understood today. To begin to provide insights, here we characterized both spatially and temporally the microbial and viral communities present in the wastewater treatment plant impacted River Erpe. We showed that within this human-impacted river, compartments were distinct in terms of chemical dynamics and microbiome composition. Interestingly, the viral communities recovered from the River Erpe overlapped more with viral communities from other wastewater treated samples than oceanic systems. Leveraging our genome-resolved databases, we were able to designate virus-host linkages for 30% of our dereplicated MAGs and show that viral predation could potentially impact several key metabolic processes from dominant microbial members that could alter carbon and nitrogen resource pools. In addition to predation, vMAGs also encoded for several auxiliary metabolic genes that may alter inorganic nitrogen availability, and possibly complement host ribosomal genes. We also identified groups of vMAGs and MAGs that were differentially stable across river compartments, and showed that on average, microbiomes in the SW are more stable than those in PW. Finally, using WGCNA networks, we identify ecologically co-occurring communities through time and show these genomes were highly related to ecosystem biogeochemistry, potentially resulting from microbial metabolisms and viral controls. Together, our results highlight the power of temporally resolved metagenomics in understanding river dynamics. Further, we provide methods and analyses that can be implemented across temporal datasets to reveal meaningful ecological patterns. Finally, this research also provides a strong scaffolding foundation for future temporally resolved river studies that leverage tools like metatranscriptomics, metaproteomics, and biogeochemical rates in order to bridge the gap between the potential impacts of river microbiomes on biogeochemistry, and their direct, quantifiable effects.

## Funding

JRR, AO, MAB, RMF, RAD, JE, LS, and MS were fully or partially supported by awards to KCW, including those from DOE Office of Science, Office of Biological and Environmental Research (BER), grant nos. DE-SC0021350 and DE-SC0023084, as well as the National Science Foundation, grant no. 2149506. A portion of this work was performed by MAB, RD, AEG, and JCS at Pacific Northwest National Laboratory (PNNL) and funded by the U.S. Department of Energy, Office of Science, Office of Biological and Environmental Research, and Environmental System Science (ESS) Program. This contribution originates from the River Corridor Scientific Focus Area (SFA) project at Pacific Northwest National Laboratory (PNNL). PNNL is operated by Battelle Memorial Institute for the U.S. Department of Energy under Contract No. DE-AC05-76RL01830. A subcontract to KCW from the River Corridor SFA also supported a portion of this work. Metagenomic sequencing was performed by the University of Colorado Anschutz’s Genomics Shared Resource supported by the Cancer Center Support Grant (P30CA046934).

## Acknowledgements

Data used in this manuscript were collected as a part of the WHONDRS 2018 sampling campaign and we thank those that participated in the design and implementation of that effort. Samples were sequenced and processed as a part of the Genome Resolved Open Watersheds effort to sequence world rivers. We also thank Tyson Claffey and Richard Wolfe for Colorado State University server management.

## Supplementary Figures and Tables

***Supplementary*** Figure 1: A) Viral accumulation curve for identified viral genomes across both the surface (SW) and pore (PW) water compartments. B) Species accumulation curve for identified microbial genomes across SW and PW compartments. C) Total richness for vMAGs across SW and PW compartments. D) Total richness for MAGs across SW and PW compartments. E) Shannon’s H for vMAGs across SW and PW compartments. F) Shannon’s H for MAGs across SW and PW compartments.

***Supplementary*** Figure 2: Procrustes analysis of the vMAG and MAG non-metric multidimensional scaling (NMDS) ordinations. Figures show the PROCRUSTES results indicating high correlation between the two communities.

***Supplementary*** Figure 3: The total proportion of clustered vMAGs that were added to the vContact2 protein clustering network. Venn diagram shows the number of vMAGs in each “group” across the different environments. Bar graphs below show the total number of vMAGs that clustered with Erpe vMAGs across ecosystems (left), and the total number of vMAGs that were used as input (right). Note, due to vContact2 quality control parameters, not all vMAGs used in protein clustering stage are retained, hence the discrepancy between total number of genomes mined from public datasets and total values in the diagram.

***Supplementary*** Figure 4: WGCNA networks of the surface water and pore water microbial and viral communities. Each circle represents a node (i.e., individual vMAG/MAG), and each line represent an edge which denotes protein cluster similarity. Modules with organisms that are predictive of an environmental variable are denoted by a yellow star.

***Supplemental Table 1****: Metadata;* sequencing_info, ESS-dive_metadata, all_geochem and subset of Geochem used for ordinations

***Supplemental Table 2****: vMAGs;* all vMAG info including genome statistics, vContact2 protein clustering results, read mapping results, annotations, AMG information, and virus-host linking results

***Supplemental Table 3****: MAGs;* all MAG info including genome statistics, taxonomy, annotation of virus-linked hosts, and mapping results.

***Supplemental Table 4****: Temporal statistics;* Results of temporal analyses i.e., bray Curtis dissimilarities, persistence, and overall genome stability.

***Supplemental Table 5****: Other statistics;* Results of other statistical analyses i.e., PERMANOVA, indicator analyses, and PROCRUSTES.

## Notes

### Competing Interest Statement

The authors have declared no competing interest.

