## Supplemental Figure 1 for "Spatial and temporal metagenomics of river compartments reveals viral community dynamics in an urban impacted stream"

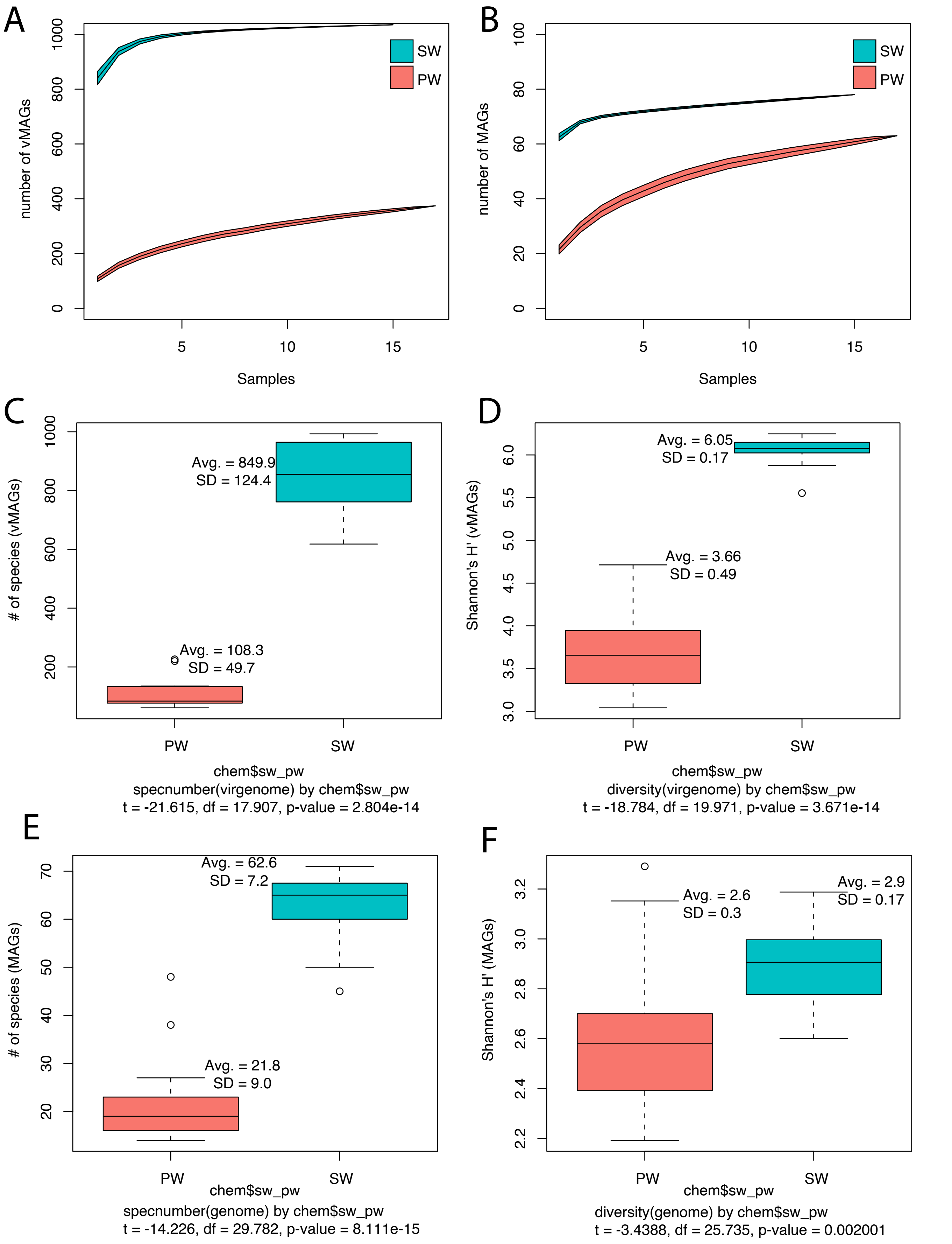


Supplementary Figure 1: A) Viral accumulation curve for identified viral genomes across both the surface (SW) and pore (PW) water compartments. B) Species accumulation curve for identified microbial genomes across SW and PW compartments. C) Total richness for vMAGs across SW and PW compartments. D) Total richness for MAGs across SW and PW compartments. E) Shannon's H for vMAGs across SW and PW compartments. F) Shannon's H for MAGs across SW and PW compartments.
