## Supplemental Figure 2 for "Spatial and temporal metagenomics of river compartments reveals viral community dynamics in an urban impacted stream"

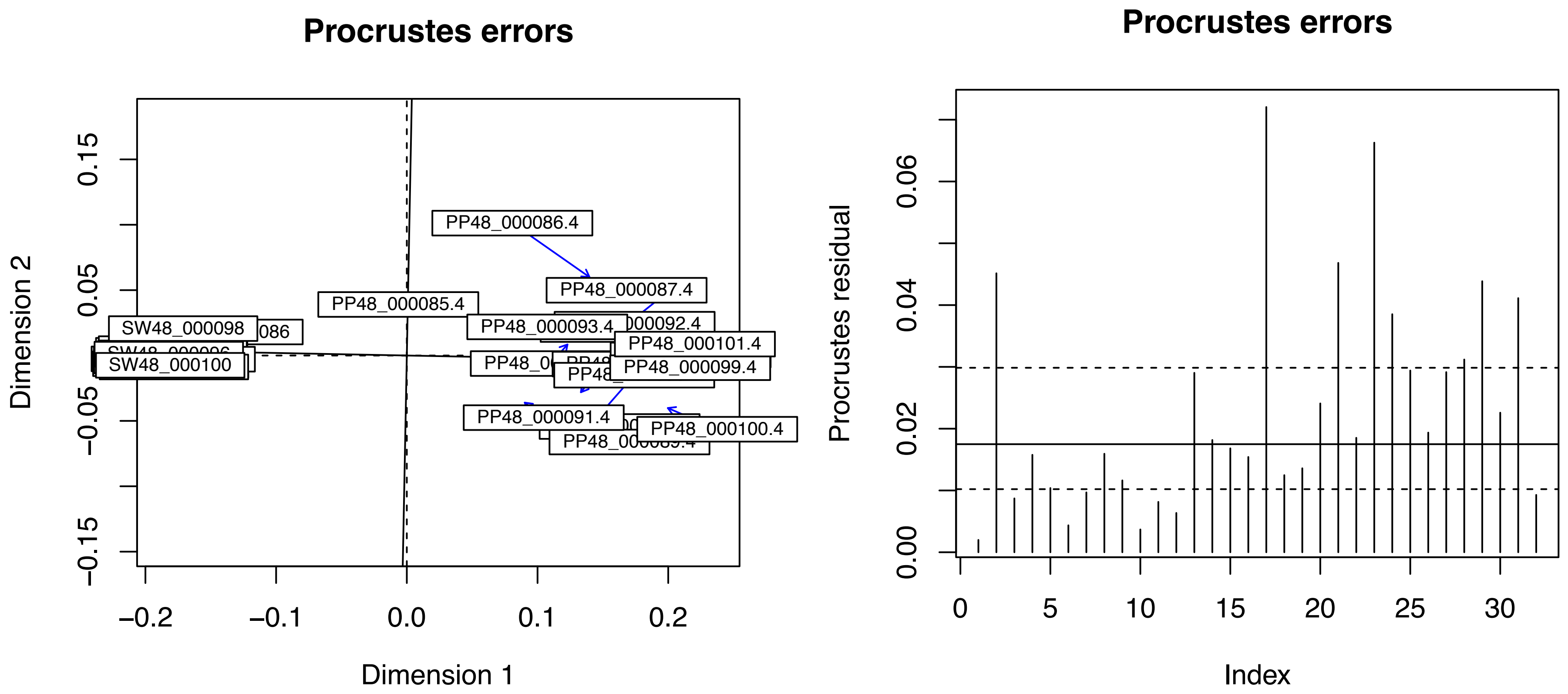


Procrustes Sum of Squares (m12 squared): 0.02662
Correlation in a symmetric Procrustes rotation: 0.9866
Significance: 0.001

Supplementary Figure 2: Procrustes analysis of the vMAG and MAG non-metric multidimensional scaling (NMDS) ordinations. Figures show the PROCRUSTES results indicating high correlation between the two communities.
