## Supplemental Figure 3 for "Spatial and temporal metagenomics of river compartments reveals viral community dynamics in an urban impacted stream"

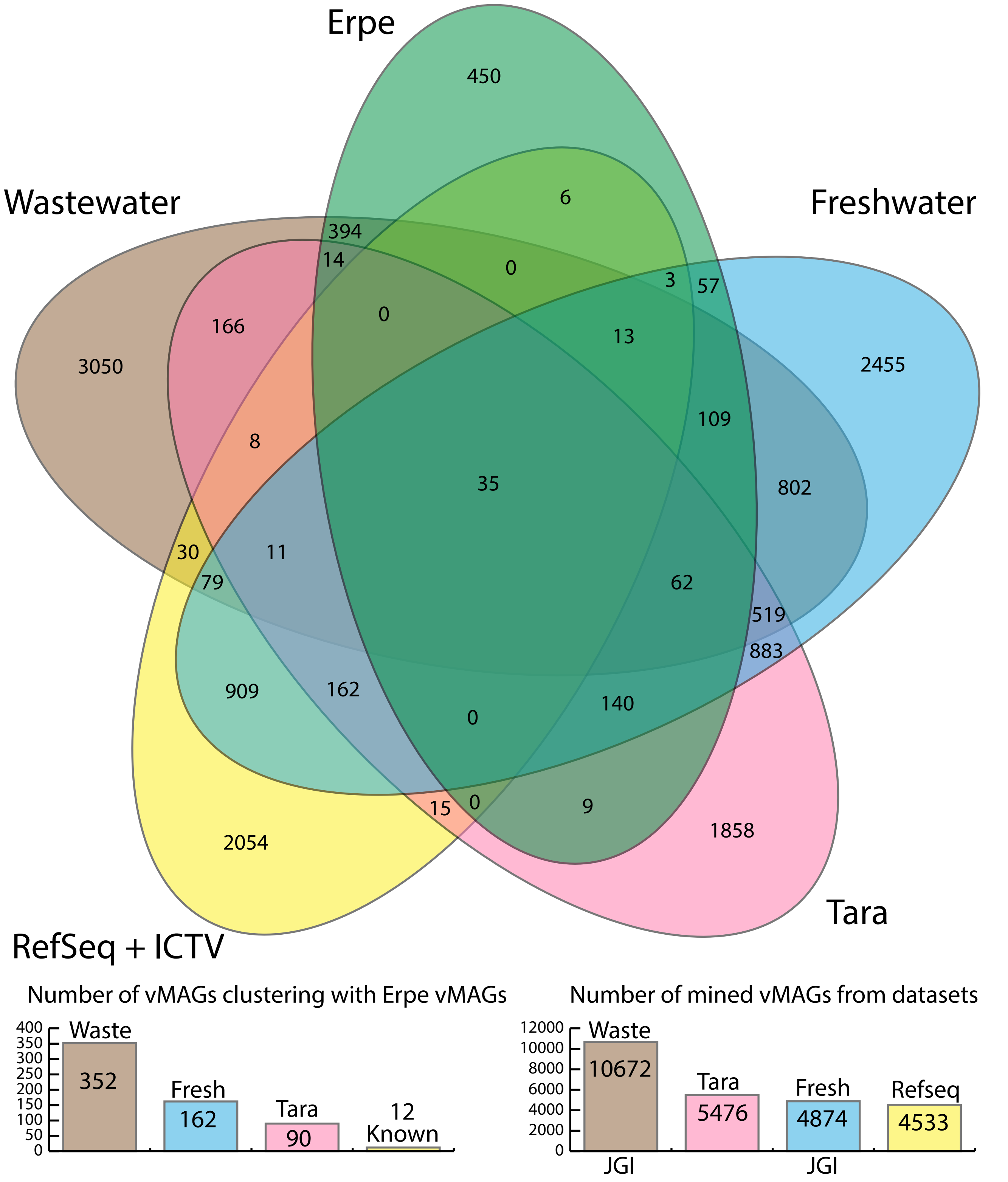


Supplementary Figure 3: The total proportion of clustered vMAGs that were added to the vContact2 protein clustering network. Venn diagram shows the number of vMAGs in each "group" across the different environments. Bar graphs below show the total number of vMAGs that clustered with Erpe vMAGs across ecosystems (left), and the total number of vMAGs that were used as input (right). Note, due to vContact2 quality control parameters, not all vMAGs used in protein clustering stage are retained, hence the discrepancy between total number of genomes mined from public datasets and total values in the diagram.
