## Supplemental Figure 4 for "Spatial and temporal metagenomics of river compartments reveals viral community dynamics in an urban impacted stream"

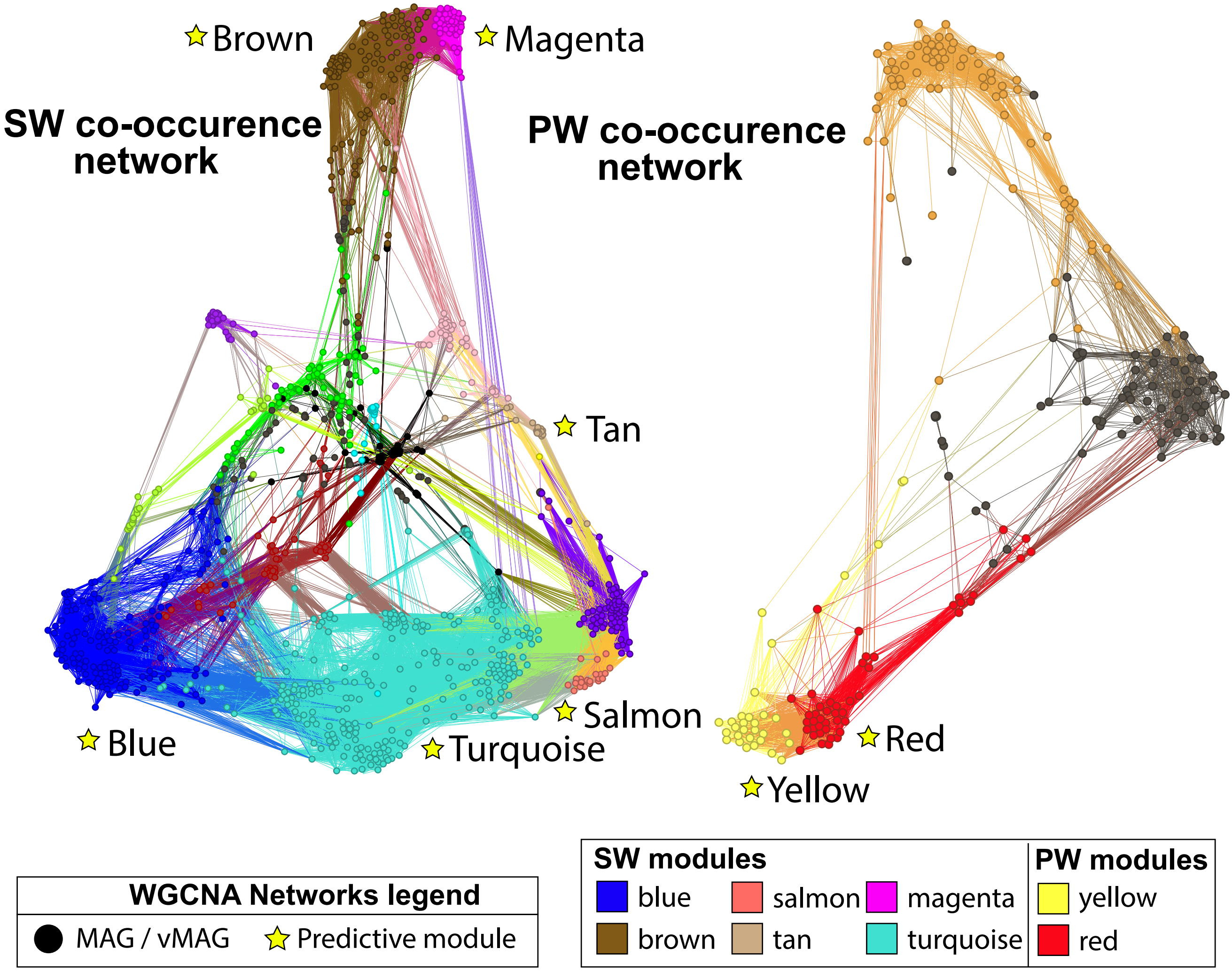


Supplementary Figure 4: WGCNA networks of the surface water and pore water microbial and viral communities. Each circle represents a node (i.e., individual vMAG/MAG), and each line represent an edge which denotes protein cluster similarity. Modules with organisms that are predictive of an environmental variable are denoted by a yellow star.
